# The ATP-bound State of the Uncoupling Protein 1 (UCP1) from Molecular Simulations

**DOI:** 10.1101/2023.03.17.533074

**Authors:** Luise Jacobsen, Laura Lydersen, Himanshu Khandelia

## Abstract

The uncoupling protein 1 (UCP1) dissipates the transmembrane (TM) proton gradient in the inner mitochondrial membrane (IMM) by leaking protons across the membrane, producing heat in the process. Such non-shivering production of heat in brown adipose tissue can combat obesity-related diseases. UCP1 associated proton leak is activated by free fatty acids and inhibited by purine nucleotides. The mechanism of proton leak remains unknown, in part due to the unavailability of high-resolution structures of the protein. As a result, the binding site of the activators (fatty acids) and inhibitors (nucleotides) is unknown. Using molecular dynamics simulations, we generate a conformational ensemble of UCP1. Using Metadynamics-based free energy calculations, we converge on the most likely ATP-bound conformation of UCP1. Our conformational ensemble provides a molecular basis of a breadth of prior biochemical data available for UCP1. Based on the simulations, we make the following testable predictions about the mechanisms of activation of proton leak and proton leak inhibition by ATP: (1) R277 plays the dual role of stabilising ATP at the binding site for inhibition, and acting as a proton surrogate for D28 in the absence of a proton during proton transport (2) the binding of ATP to UCP1 is mediated by residues R84, R92, R183, and S88 (3) R92 shuttles ATP from the E191-R92 gate in the inter-membrane space to the nucleotide binding site, and serves to increase ATP affinity (4) ATP can inhibit proton leak by controlling the ionisation states of matrix facing lysine residues such as K269 and K56 and (5) fatty acids can bind to UCP1 from the IMM either via the cavity between TM1 and TM2 or between TM5 and TM6. Our simulations set the platform for future investigations into the proton transport and inhibition mechanisms of UCP1.

## 1 Introduction

Control of excessive body fat and management of obesity-related diseases is among the most significant global challenges. Recent evidence establishes a negative correlation between the expression of brown adipose tissue (BAT) and occurrence of diabetes, coronary artery disease and hypertension in humans.^1^ Activation and increased expression of BAT is therefore a widely explored strategy for countering the obesity pandemic. BAT activation requires acute cold exposure, which is an unviable long-term clinical strategy. BAT contains high concentrations of the uncoupling protein 1 (UCP1)^2,3^ in the inner mitochondrial membrane (IMM). Under physiological stimuli such as cold exposure, UCP1 generates heat by dissipating the mitochondrial transmembrane proton (H^+^) gradient, which otherwise powers the production of ATP from ADP by the mitochondrial ATP synthase enzyme complex: a phenomenon referred to as mitochondrial uncoupling. The process entails the conversion of potential energy stored in the membrane to heat, instead of work in the form of ATP synthesis. In this way, the organism produces heat (also called non-shivering thermogenesis), and removes fats and sugars from the blood. UCP1 is not constitutively active. One way of combating obesity-related diseases is to increase UCP1 activity on demand in obese/diseased individuals by external administration of drugs. Dinitrophenol (DNP) is an archetypal mitochondrial uncoupler,^4^ but has severe side effects. BAM15 ^5^ and OPC-163493^6^ are two uncouplers developed recently which have anti-diabetic effects in rats, but are not yet approved in the clinic. Notwithstanding these scattered attempts, the development of UCP1-targeting therapy is mainly hampered by our lack of comprehension of the mechanism of proton uncoupling mediated by UCP1. Although UCP1 was discovered more than 40 years ago^2,3^ its molecular mechanism of function is still unknown. In this work, we present the first known computationally-derived conformational ensemble of UCP1, which will drive significant progress towards a mechanism of UCP1-mediated H^+^ transport.

Mg^2+^-free^7^ purine di- and triphosphates (GDP, GTP, ADP and ATP) inhibit UCP1 H^+^ transport by binding from the intermembrane space (IMS) to three arginine residues R84, R183, and R277 in the central cavity of UCP1.^8–10^ However, the precise binding configuration of ATP and the exact mechanism of inhibition is unknown. Inhibition by ATP is strongly pH dependent and decreases drastically as pH increases above 6.3.^7,11,12^ At high pH, a salt bridge between anionic E191 (E191^-^) and R92 is hypothesized to obstruct the entrance to the ATP binding site.^10,13,14^ At low pH the salt bridge is broken when E191 is protonated (E191p).^10,13,14^ Since we use ATP as the model purine nucleotide, we sometimes refer to the nucleotide binding site as the ATP-binding site in the remainder of the article.

UCP1 mediated H^+^ transport is activated by fatty acids in the IMM. In the so-called cycling model, fatty acids cycle back and forth between the leaflets of the IMM while binding protons from the IMS and releasing them to the matrix. In other models, a fatty acid acts as a temporary proton acceptor in a series of proton-accepting and releasing chemical moeities which allows H^+^ passage through UCP1 (Fig. S1). These models are referred to as the buffering ^15,16^ and shuttling models.^17^ The precise role of fatty acid and its interaction with UCP1 during H^+^ transport is still under debate, but the fatty acid recycling mechanism is mostly debunked, based on recent patch-clamp measurements on mitocytes. ^15,17^

The binding of cardiolipin lipids to UCP1 stabilises the protein^18,19^ and increases UCP1’s ability to transport H^+^ in liposomes.^18^ The crystal structure of the ADP/ATP carrier (AAC), has three bound cardiolipins. ^20,21^ UCP1 belongs to the same family of carriers and mass spectrometry showed that UCP1 also has three bound cardiolipins. ^19^

Here we perform all-atom and coarse-grained molecular dynamics (MD) simulations of UCP1 in a realistic IMM environment to realise a plausible apo-, ATP-bound and fattyacid bound conformational ensemble for UCP1. The simulations provide strong evidence in favour of the proposed E191-R92 salt bridge that gates the ATP binding site. We simulated several binding events of ATP at the ATP binding site using standard all-atom simulations and by Multiple Walker Well Tempered Metadynamics simulations to evaluate the lowest free energy configurations of bound ATP. We found that the presence of negatively charged ATP in the IMS facing cavity of UCP1 increases the *pKa* of several lysine residues on the matrix facing side of UCP1. Such transitions of ionisation states are indicative for a potential inhibition mechanism. Finally, we find that anionic, but not neutral fatty acids can bind to UCP1 via the TM1/TM2 or the TM5/TM6 interface.

## 2 Results and Discussion

### The UCP1 Model is Impermeable to Water

No experimental structure of UCP1 existed at the time of writing this manuscript. Prior investigations of UCP1 relied on homology models based upon the AAC structure template.^22^ While the secondary structures of AAC and UCP1 are highly similar^23^ their amino acid sequences have only 41% similarity (23% identity), which can be inadequate for AAC to be a reliable homology model template. An alternative template is the structure of the uncoupling protein 2 (UCP2, pdb: 2lck)^24^ which has 74% similarity (58% identity) to UCP1. However, it was previously demonstrated ^25^ that the 2lck structure obtained from NMR in detergents does not reflect a legitimate UCP2 conformation. For example, in the 2lck structure, there is a continuous water channel connecting the matrix and IMS (Fig. 1B). Water can directly cross the IMM through this channel carrying protons with it all the time, leading to uncontrolled dissipation of the proton gradient.

**Figure 1:**
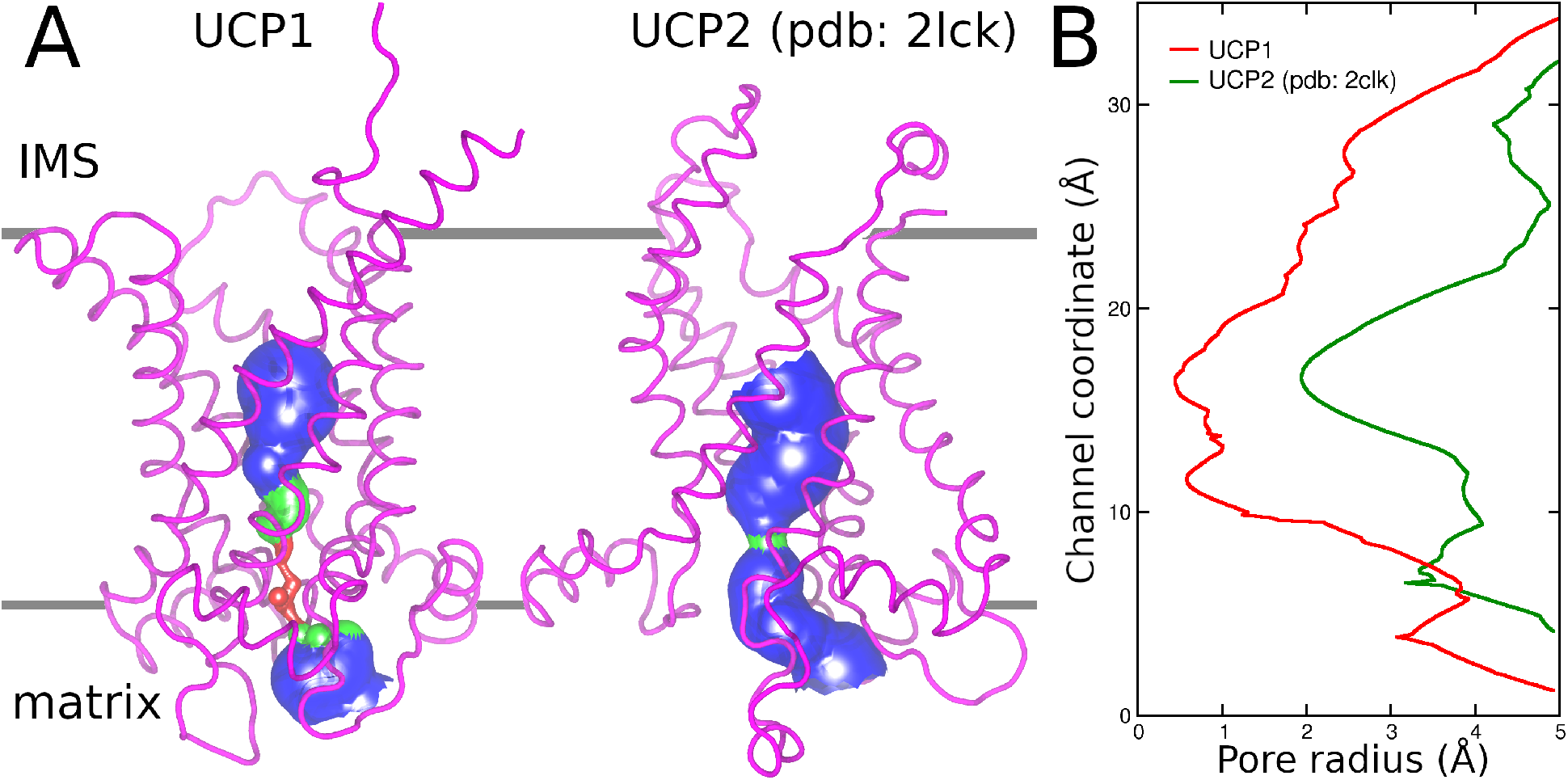
**A**: The radius of the cavity through the UCP1 AlphaFold model after 500 ns of simulation as compared to the experimental structure of uncoupling protein 2 (UCP2, pdb: 2lck).^24^ The proteins are illustrated as purple tubes and the surface indicates whether the cavity is solvent accessible (blue), can fit one water molecule (green), or is solvent inaccessible (red).^26^ **B**: Pore radius along the protein cavity in the UCP1 model (red) and experimental UCP2 structure (green).

Here, we use all-atom MD simulations of the UCP1 AlphaFold^27,28^ model (5 replicas of 500 ns each) to derive a reliable structural ensemble of UCP1. The AlphaFold UCP1 model possesses the three fold symmetry typical of the SLC25 family of mitochondrial carriers of which UCP1 is a member^29^ (Fig. S2). UCP1 folds into three repeated domains of ~100 residues that each contains two transmembrane *α*-helices (H1-H6), which fold into a helix pair, and a matrix facing short *α*-helix and loop connecting to the neighbouring repeat.^23^ In contrast to the UCP2 structure (pdb: 2lck), the AlphFold UCP1 model is impervious to water, with no continuous conduit connecting the matrix and IMS. UCP1 remains in this impervious configuration throughout simulations in an IMM environment (Fig. 1).

### Three Cardiolipin Lipids Bind to UCP1

Cardiolipin lipids directly bind and stabilize UCP1, and increase H^+^ transport across the IMM in liposomes.^18,19^ In total, three cardiolipins are predicted to bind UCP1 in the same manner as to the AAC (pdb: 2C3E and 1OKC),^20,21^ i.e., to the [YF]xG and [YWF][KR]G motifs at the negative end of the helix dipoles of the matrix helices and the even-numbered transmembrane helices (Fig. 2A). ^19^

**Figure 2:**
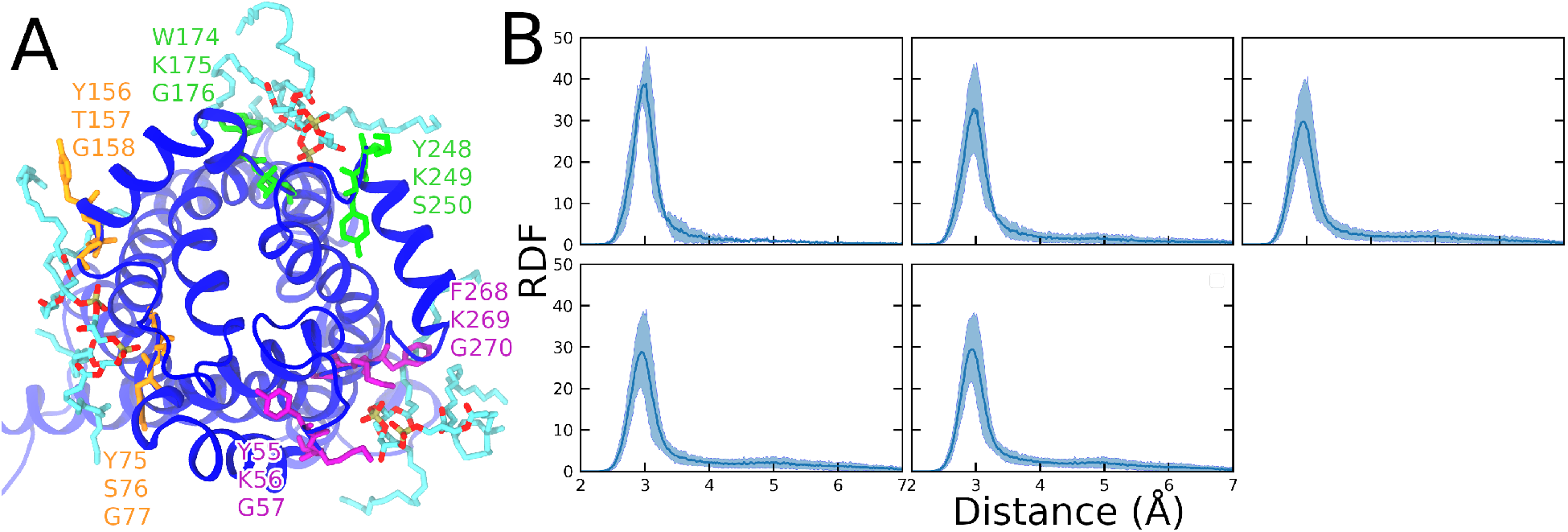
**A**: UCP1 (blue cartoon) seen from the matrix with three cardiolipins (turquoise (C), red (O), and brown (P) colored stick) bound to the predicted [YF]xG and [YWF][KR]G motifs (orange, purple, and green stick).^19^ **B**: Radial distribution function (RDF) of phosphate atoms in the three bound cardiolipins with respect to UCP1 in five independent simulations of unbound UCP1 in an IMM. The RDFs are averaged over the three cardiolipins and 500 ns of simulation.

In three 3 μs coarse-grained MD simulations of UCP1 with lipids randomly distributed in the membrane plane (see Table 3 for membrane composition), cardiolipin lipids spontaneously diffuse to and bind UCP1 at the three predicted cardiolipin binding sites (Fig. S3). The simulations, together with the previous experimental evidence, show that the most natural state of UCP1 is with three cardiolipin lipids bound. Therefore, in the all-atom simulations, cardiolipins were pre-positioned at the three cardiolipin binding sites where they remained bound throughout all the performed simulations (Figs. 2B, S3-S6).

**Table 1:** Mean and standard deviations of *pKa* of matrix facing lysine residues whose average *pKa* is below the pH of the matrix solvent (7.8) in the simulations of unbound UCP1.

| residue | unbound UCP1 (E191<sup>-</sup>) | UCP w. ATP (E191<sup>-</sup>) |
| --- | --- | --- |
| **K56** | 7.25 ± 0.71 | 9.59 ± 0.88 |
| **K67** | 7.45 ± 0.65 | 10.06 ± 0.71 |
| **K73** | 7.53 ± 0.65 | 9.59 ± 0.72 |
| **K269** | 7.29 ± 0.87 | 9.30 ± 0.93 |

**Table 2:** Average hydrogen bond occupancy between anionic fatty acids and Arginine residues of UCP1 which had average occupancies above 15% for at least two fatty acids species during the ten 500 ns simulations of lauric (C12:0), palmitic (C16:0), palmitoleic (C16:1), stearic (C18:0), or linoleic (C18:2) acid.

|  | **R71 (%)** | **R162 (%)** | **R294 (%)** | **R300 (%)** |
| --- | --- | --- | --- | --- |
| **C12:0** | 29.3 | 8.0 | 23.0 | 20.6 |
| **C16:0** | 6.7 | 15.5 | 0.0 | 9.2 |
| **C16:1** | 11.6 | 42.9 | 19.7 | 33.5 |
| **C18:0** | 40.0 | 3.7 | 23.4 | 9.2 |
| **C18:2** | 44.9 | 12 | 13.4 | 0.1 |

**Table 3:**
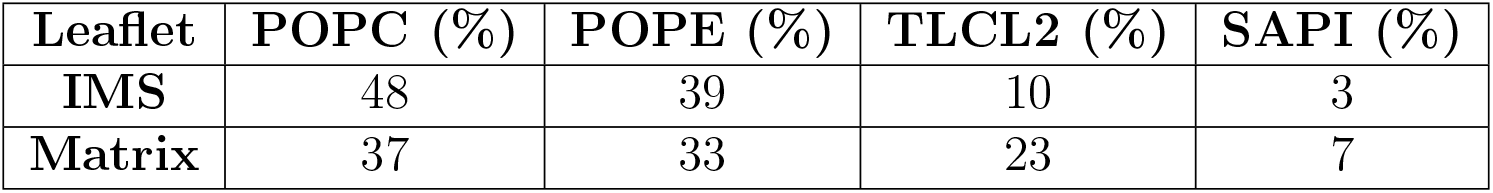
Membrane composition used in the simulations to mimic the IMM. POPC: 3-palmitoyl-2-oleoyl-glycero-1-phosphatidylcholine, POPE: 3-palmitoyl-2-oleoyl-glycero-1-phosphatidylethanolamine, TLCL2: tetralinoleoyl cardiolipin (head group charge = −2), SAPI: 3-stearoyl-2-arachidonoyl-glycero-1-phosphoinositol. The lipid definitions follow CHARMM-GUI.

In five replicas of 500 ns all-atom simulations of ligand-free UCP1 in an IMM, UCP1 (1) retained the expected secondary structure, (2) was impervious to water between the matrix and IMS, (3) had three cardiolipins bound at the predicted binding sites, and (4) had a backbone root mean squared deviation (RMSD) that converged (Fig. S7). Therefore, we concluded that the conformational ensemble of apo-UCP1 is reliable and ready to be used to investigate the interaction of UCP1 with ATP and fatty acids.

### Access to the ATP Binding Site is Gated by a Salt Bridge

To probe an ATP bound configuration of UCP1, ten all-atom MD simulations with ATP initially placed at the entrance to the ATP binding site were launched for ~450 ns each. ATP was placed with different initial orientations near the entrance. In five simulation replicas (Rep1-Rep5), the nucleobase pointed towards the binding site. In the five other simulations (Rep6-Rep10) the triphosphate group pointed towards the binding site (Fig. S15).

ATP did not approach the binding site in any of the ten simulations (top panel of Fig. 3D and S15). In four simulations, a salt bridge between E191^-^ and R92 obstructed access to the ATP binding site, as predicted by Klingenberg and co-workers, ^7,10,12^ (Fig. 3A, C, and E top panel). In simulations of ligand-free UCP1, the average *pKa* of E191 across all five replicas was 7 (Fig. S16). E191 is likely to be protonated because it faces the IMS (pH ~6.8). Therefore, all 10 simulations with ATP were repeated but with E191 protonated (E191p).

**Figure 3:**
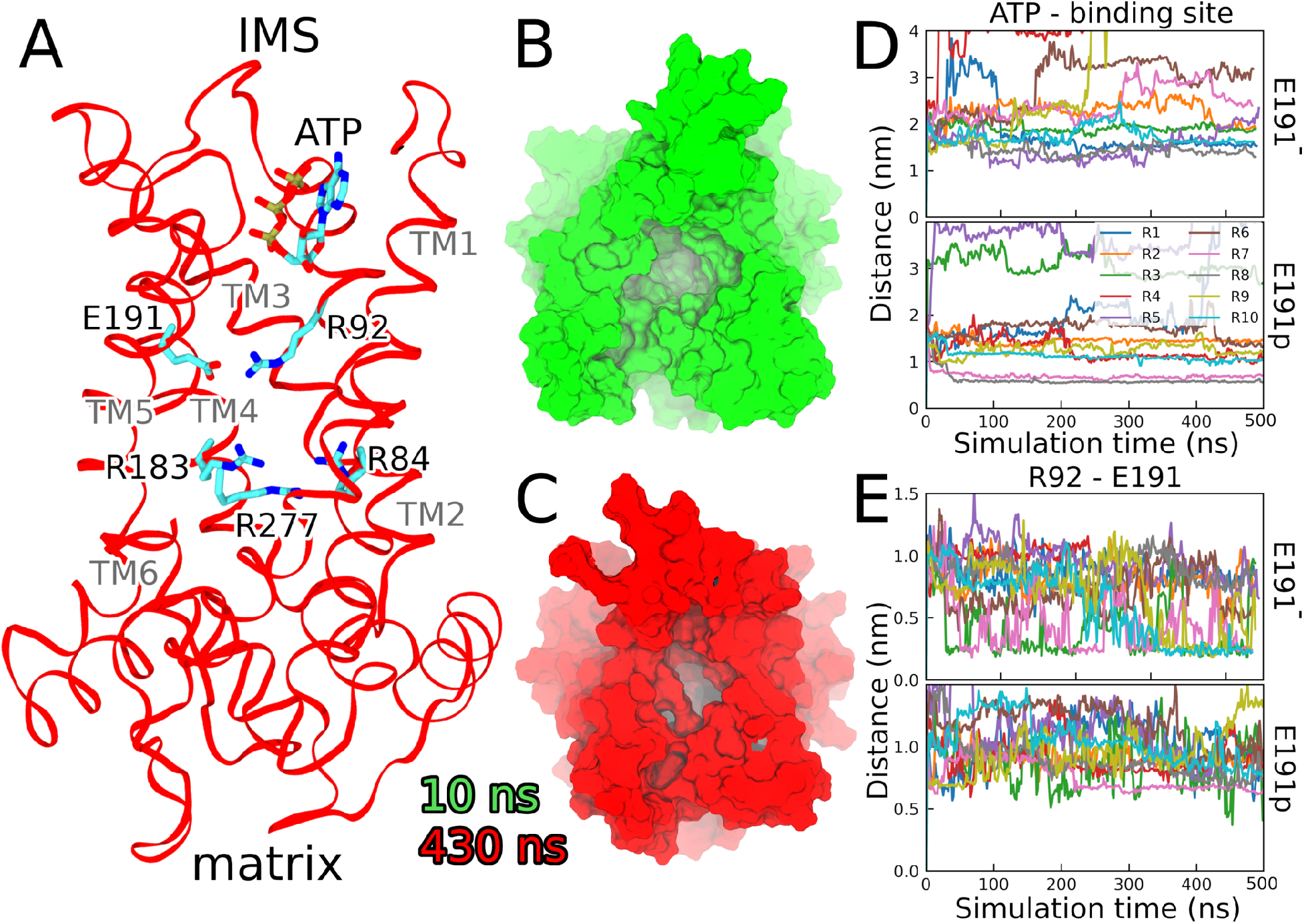
**A**: UCP1 with E191^-^, the salt bridge (E191 and R92), and the ATP binding site (R84, R183, and R277) after 430 ns. Most of TM6 is omitted to improve visibility. Surface representation of UCP1 after **B**: 10 ns (green) and **C**: 430 ns (red). **A**-**C** are from replica Rep3. **D**: Distance between the centre of mass of the ATP molecule and the ATP biding site (R84, R183, and R277) in simulations with E191^-^ (top) and E191p (bottom). **E**: Distance between the E191 carboxyl oxygen and the R92 guanidinium nitrogen in simulations with E191^-^ (top) and E191p (bottom). R1 through R10 refer to replicas Rep1 through Rep10.

In the simulations with E191p, the salt bridge no longer formed (Fig. 3E bottom panel) which allowed ATP to reach the ATP-binding site in simulation replicas Rep7, Rep8, and Rep10 (Fig. 3D bottom panel). In this way, protonation of E191 and the opening of the E191-R92 gate is an essential step in the binding of ATP and ATP-induced inhibition of H^+^ transport. The ATP molecules that entered the binding site (in simulations Rep7, Rep8, and Rep10) started with the triphosphate group pointing towards the binding site. The three ATP molecules that drifted furthest away from the binding site (Rep1, Rep3, and Rep5) initially had the nucleobase pointing towards the binding site. Therefore, we conclude that ATP binding and inhibition is initiated by ATP entering the binding site with the triphosphate group first. These observations are somewhat different from simulations of ADP binding to the AAC, where ADP bound within a few nanoseconds to the ADP-binding site irrespective of the initial ADP orientation. ^30^ The differences arise from the initial placement of the nucleotides and the differences in the nucleotide binding sites of AAC and UCP1. In the simulations of AAC, ADP was placed closer to the vestibule of AAC than we place ATP. Secondly, the AAC nucleotide binding site contains more cationic amino acids. ^31^

### R92 Shuttles ATP to the Binding Site

If R92 was only involved in the salt bridge with E191, and played no further part in ATP binding, then replacement of R92 by an uncharged residue would not influence the binding affinity of ATP to UCP1. However, the R92T mutation maintains ATP binding to UCP1, but with a significantly lower binding affinity and rate than in wild-type UCP1.^10^ Therefore, R92 directly influences the binding affinity. The R92T mutant also maintains a significant ATP inhibition of fatty acid transport, unlike mutations in the other arginine residues: R84, R183 and R277. ^10^ The simulation data explain this dual role of R92. In all the three binding events in the simulations, we find thawt R92 follows the ATP molecule and stabilises it at the binding site (Figs. 4 and S20). R92 is flexible within the cavity of UCP1 and can readily move its guanidium group from the IMS, where it binds ATP, to the ATP binding site as can be seen during the binding event of ATP in simulation Rep8 (E191p) (Video ATP_binding.mov). We propose that that R92 acts as a shuttle which aids the transport of ATP from the IMS to the ATP binding site.

**Figure 4:**
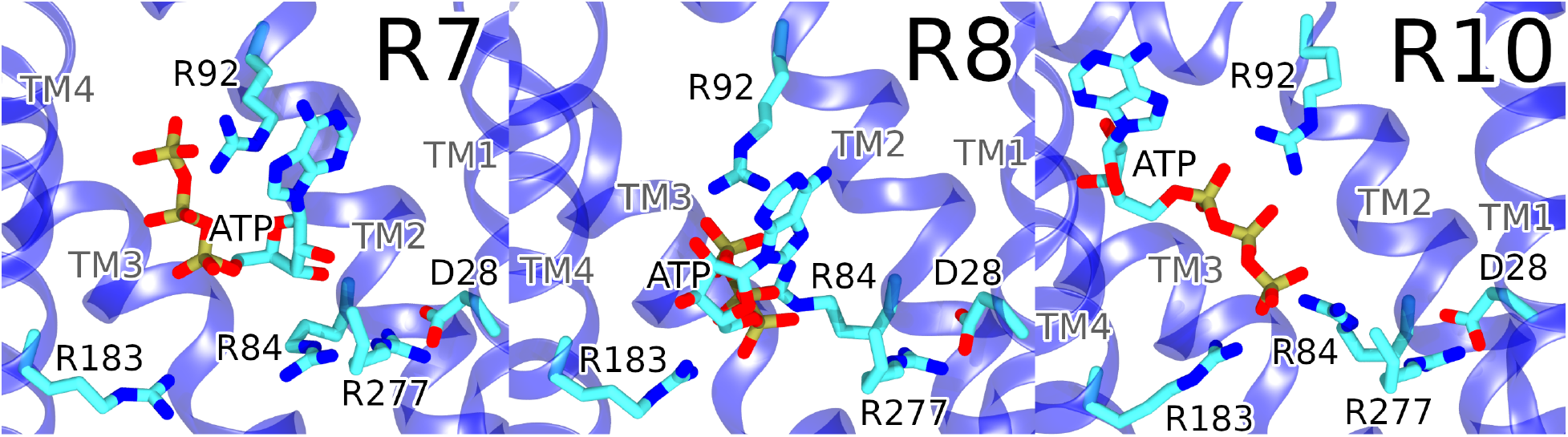
Final binding configurations of ATP in simulations Rep7, Rep8, and Rep10 with E191p. Panels R7, R8 and R10 refer to replicas Rep7, Rep8 and Rep10

### The ATP Binding Site

None of the three obtained ATP-bound configurations obtained from the aforementioned simulations were alike (Fig. 4). In Rep7, ATP only bound R92 directly. In Rep8 the triphosphate group bound R92, R84, and R183. The same was the case in Rep10, except here the ATP molecule adopts a more elongated configuration in the cavity and its center of mass was further from the ATP binding site than in Rep8 (Figs. 3 and 4). R277 interacted with D28 in all the three simulations (Fig. S21), but did not directly bind ATP. The protonation-mimicking D28N mutation drastically reduces H^+^ transport.^32,33^ Protonation of D28 is likely to break the salt bridge between R277 and D28. Modriansky et al., proposed that R277 was essential for ATP inhibition of UCP1, but not required for binding.^9^ In other words, inhibition will require that binding of ATP induces a conformational change of UCP1 leading to a tight binding state which involves binding to R84, R183, and R277.^9^ The binding configurations we obtain possibly represent the initial binding of ATP to the binding site and do not involve direct binding to R277. It is worth noting that in contrast to Modriansky et al., Echtay et al. ^10^ demonstrated that ATP binding was completely dependent on R277. Although R277 does not directly coordinate ATP in our simulations, it can certainly contribute to the overall positive charge density in the binding pocket and thereby influence the binding affinity of ATP.

D28 most likely acts as one of several proton acceptors in the IMS-open state of UCP1. In the absence of a H^+^ or ATP, the negative charge on D28 is compensated by a salt bridge made with R277, reminiscent of the K791 residue in the gastric proton pump, which acts as a proton surrogate in conformations of the cycle when the pump is not bound to protons. ^34^ In this way, R277 supports H^+^ transport. In the presence of ATP, however, R277 serves to increase the binding affinity of ATP. In this way, R277 plays a dual role in UCP1: it can contribute both to the inhibition of UCP1 in the presence of ATP, and contribute to H^+^ transport by serving as an intermittent proton surrogate.

To examine which binding configuration (Rep7, Rep8, or Rep10) was most likely, we initiated Multiple Walker Well Tempered Metadynamics simulations^35,36^ from the three configurations both with and without a transmembrane electric potential of ~160 mV. The RMSD of the ATP molecules with respect to the initial binding conformation of Rep8 (Fig. 4) was chosen as the collective variable. The metadynamics simulations measure the relative free energy between the different binding configurations. Metadynamics simulations were performed both with and without a transmembrane electrical potential.

With or without a transmembrane electric potential, the ATP molecule in Rep7 drifts away from the protein within the first 1 μs (Fig. 5). Therefore, ATP binding to only R92 is weaker than binding which also involves R183 and R84. The ATP molecule in Rep10 stays UCP1-bound. However, the RMSD of ATP in Rep10 fluctuates significantly, both in simulations with or without a transmembrane electric potential, suggesting that the ATP molecule is not confined to a single binding configuration. On the contrary, there are only small fluctuations in the RMSD of the ATP molecule in Rep8 (Fig. 5A,B). Additionally, the RMSD in Rep8 stays within 0-0.7 nm, which demonstrates that ATP binding is more stable in Rep8 than in Rep7 and Rep10.

**Figure 5:**
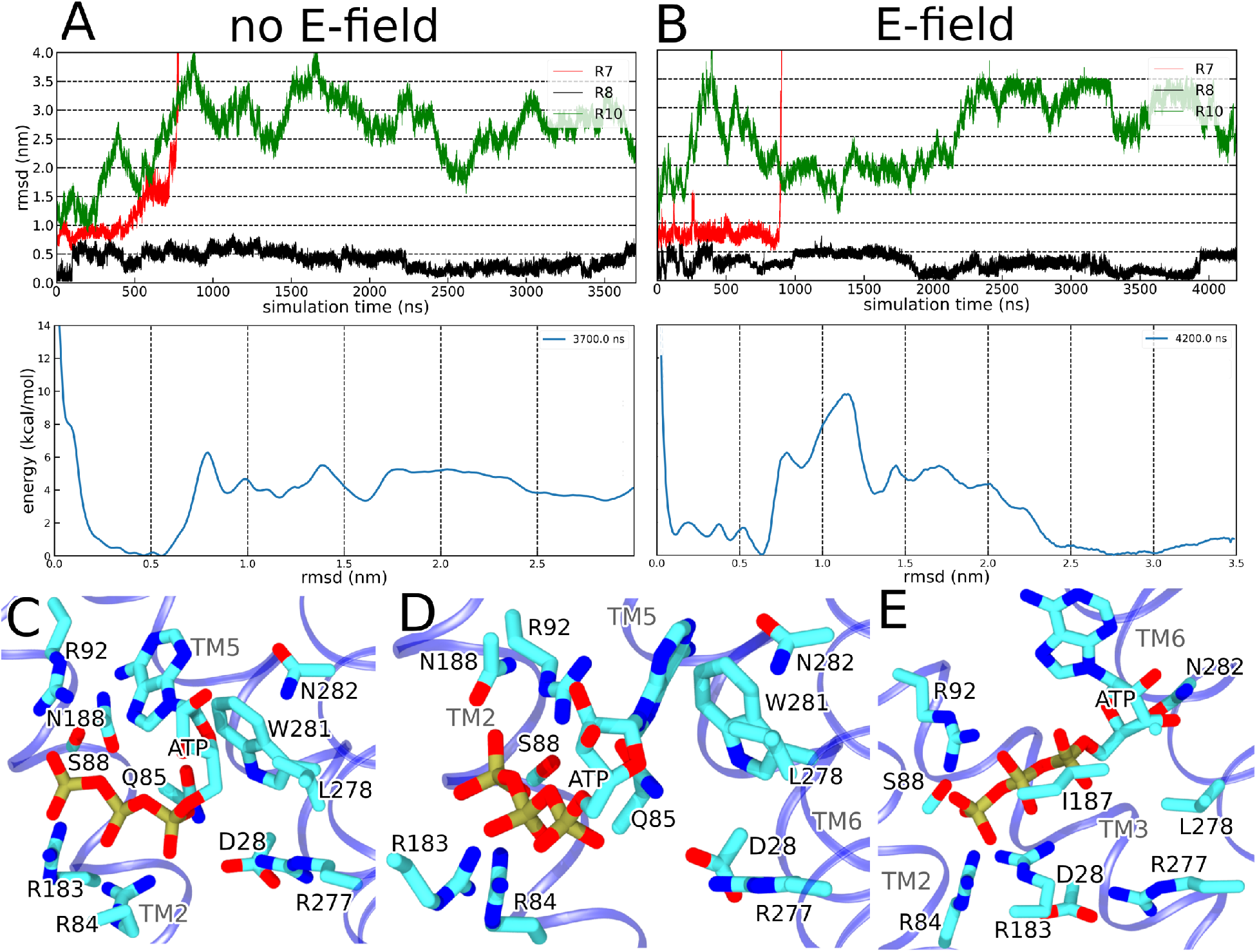
Evolution of the RMSD of ATP as a function of simulation time (top panel) and the energy landscape (mid panel) obtained from Multiple Walker Metadynamics simulations continued from simulations Rep7, Rep8, and Rep10 without (**A**) or with (**B**) a transmembrane electric potential (~160 mV). ATP binding configurations in Rep8 from metadynamics simulations without (**C-D**) and with (**E**) a transmembrane potential. In the top panels, R7, R8 and R10 refer to replicas Rep7, Rep8 and Rep10

With or without a transmembrane potential, the minimum in the free energy profile extracted from Metadynamics hovers between an RMSD of 0.4 and 0.7 nm corresponding to the binding conformations sampled in the Rep8 system (Fig. 5A, B). Hence, the Rep8 configuration is the most likely ATP-bound conformation of UCP1. A second minimum appears around 2.8 nm in the energy landscape with a membrane potential. The minimum at 2.8 nm is wide and arises from configurations sampled in Rep10, which are much less stable than in Rep8. Therefore, the minimum at 2.8 nm is not considered a promising candidate to represent a reliable ATP-bound configuration. It must be noted that despite application of the same bias on all three conformations, the collective variable is not fully sampled, and therefore the free energy profile may not be converged (Fig. S22 and S23). Based on the current set of simulations, however, the Rep8 configuration is most likely.

Three slightly different ATP-bound conformations emerge from Rep8: the original conformation (0-0.4 nm, Fig. 5D) and one at 0.4-0.7 nm which is different for the simulation without (Fig. 5C) and with (Fig. 5E) a membrane potential. Common for the three binding conformations are continuous interactions with R84, R183, S88, and L278. R84, R183, and S88 accommodate the charged triphosphate group while L278 interacts with the ribose sugar. Only the orientation of the nucleobase differs noticeably between the binding conformations in Rep8. The conformation at 0.4-0.7 nm in the simulation without a membrane potential (Fig. 5C) has the nucleobase tilted towards N282 and W281 with the latter participating in aromatic stacking with the nucleobase. Meanwhile, the binding configuration around 0.5 nm in the simulation with a membrane potential (Fig. 5E) has the adenosine group tilted into the centre of the protein with no direct protein interaction.

Figure 6 compares the nucleotide-bound conformations of UCP1 and AAC. No structure of ADP-bound AAC is available. Therefore, we ran simulations of the ADP-bound configurations of AAC with the same protocol as described for UCP1. In AAC, the binding of nucleotides is highly specific, ^37^ because the binding free energy is likely used in the conformational change required for the exchange of ADP and ATP. However, in UCP1 the binding site is more promiscuous and accommodates several nucleotides.^38–40^ Residues R84, R183 and S88 of UCP1 accommodate the charged phosphate moieties of ATP. By comparison, in AAC, the phosphate moieties of ADP bind to R79 (R84 in UCP1), K22 (D28 in UCP1) and R279 (R277 in UCP1). Whereas UCP1 shows limited coordination of the nucleotide base, simulations of AAC reveal a stacking of the adenine moiety with Y186, which pulls the adenine to a pocket consisting of I183, G182 and S227. This adenine-binding pocket is well conserved within the carrier sub families and differentiates according to the translocated species.^41^ However, in UCP1, no such adaptation for the binding of nucleotide bases is seen, allowing for promiscuity towards both guanine and adenine nucleotide binding.

**Figure 6:**
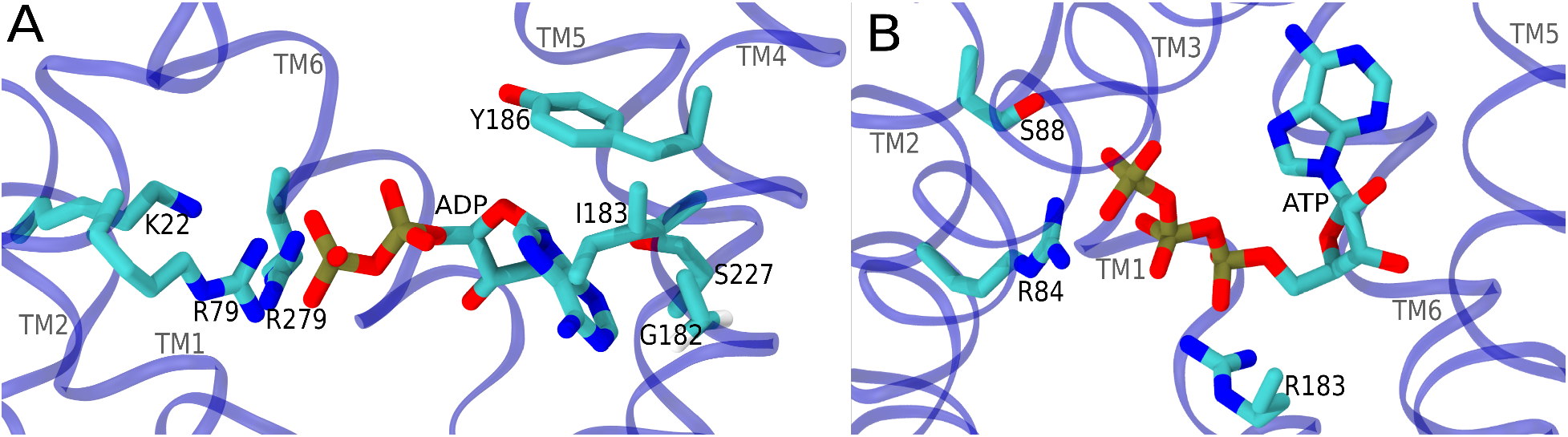
**A**: The ADP-bound configuration of AAC and **B**: ATP-bound configuration of UCP1. Both are final simulation snapshots.

### ATP Switches the pKa of Lysine Residues on the Matrix side

ATP has a charge of −4, hence entry of ATP into the binding pocket will significantly alter the electrostatic environment in the binding pocket, leading to changes in the ionisation states of amino acid residues. A change in the ionisation state of residues such as Glutamate and Aspartate can influence H^+^ transport, if the residues are protonated and deprotonated when an H^+^ is transported through UCP1. Such dynamic ionisation states are proposed to be responsible for *K*^+^ selectivity in P-type ion pumps which employ several anionic residues in the ion-binding site.^42–44^ In principle, ATP binding can inhibit proton transport simply by confining an ionisable residue to a single ionisation state.

ATP alters the *pKa* values of Lysine residues which are exposed to the matrix and distant from the ATP-binding site. The Lysine side chain has a standard *pKa* of 10.54 and changes in the ionisation state of the Lysine side chain are rarely observed.^45^ For UCP1, in the absence of ATP, K56, K67, K73, and K269 had *pKa* values below the pH (~7.8) of the surrounding solvent in the matrix (Table 1 and Fig. S17), suggesting that they are likely to be deprotonated (neutral) in ligand-free UCP1.

However, in the simulations where ATP is placed at the mouth of the ATP binding site, K56, K67, K73, and K269 all had an average *pKa* well above 7.8 and are likely to be protonated (cationic). With ATP, the *pKa* of K56, K67, K73, and K269 is either very stable around 10-11 or fluctuates rapidly between two values: one around 10 and one around 7 (Figs. S18 and S19). Note that K56, K67, K73, and K269 face the matrix, while ATP enters from the IMS. There are no direct molecular contacts between ATP and the lysine residues. Yet, ATP exerts a significant electrostatic effect on the ionisation states of Lysine residues facing the matrix. K56 and K269 are proposed to be involved in binding of fatty acid and in H^+^ transport^46^ based on NMR investigations of UCP1 in micelles. An older investigation suggested that fatty acids could activate H^+^ transport in a K269 deletion mutant (which also deleted F268 and G270), but the mutant was insensitive to inhibition by ATP.^47^

Based on the *pKa* values, it can be concluded that the binding of ATP switches K269 from a neutral to a cationic state. Once K269 is deleted, ^47^ there will never appear a positive charge in the K269-containing loop despite binding of ATP. Since ATP inhibits UCP1, the *pKa* switch implies that K269 must be neutral for effective H^+^ transport. In other words: a positive charge must not manifest in the loop containing K269. Fatty acids can still activate a K269 deletion mutant, ^47^ because such a mutant will never acquire a positive charge. In this way, the *pKa* switch of K269 (and possibly K56, K67 and K73 by a similar mechanism) may be linked to ATP inhibition. It remains to be seen how a sustained *pKa* below 7.8 supports FA-mediated activation of H^+^ transport, as opposed to a higher *pKa* acquired upon ATP binding.

### Fatty Acids Bind to UCP1 via the TM1/TM2 or the TM5/TM6 interface

To find fatty acid interaction sites in UCP1, we set up all-atom simulations of UCP1 with five different fatty acids. ATP-free UCP1 was embedded in a membrane of composition modelling the IMM (Table 3) with an additional 10% (12-14 in number) fatty acid molecules in each leaflet. The fatty acids (lauric (C12:0), palmitic (C16:0), palmitoleic (C16:1), stearic (C18:0), or linoleic (C18:2)) were distributed randomly in each leaflet and a membrane potential of ~160 mV was applied for these simulations. Half the fatty acids were neutral (protonated), while the other half were anionic. Ten 500 ns replicas for each system were simulated, resulting in a total of 25 μs of simulations, which are analysed below.

Neutral fatty acids did not have specific interactions with UCP1 (Fig. S24). Anionic fatty acids bind to UCP1 predominantly via arginine residues of UCP1. The interaction sites for anionic fatty acids are located near R71 and R162 facing the matrix and R294 and R300 facing the IMS (Fig. 7 and Table 2). R294 and R300 are both on TM6 while R71 is located on the short matrix facing helix between TM1 and TM2. R162 is on the short matrix facing helix between TM3 and TM4 (Fig. 7A). R71, R294, and R300 are all located on or near TM1 and TM6. Prior NMR experiments have shown that palmitic acid (C16:0) binds to this region. ^46^ Furthermore, D28 and R277, which are key residues in the UCP1 H^+^ transport mechanism, are located on TM1 and TM6, respectively. Thus our simulations provide evidence that a H^+^ translocation pathway through UCP1 is likely to be along TM1 and TM6.

**Figure 7:**
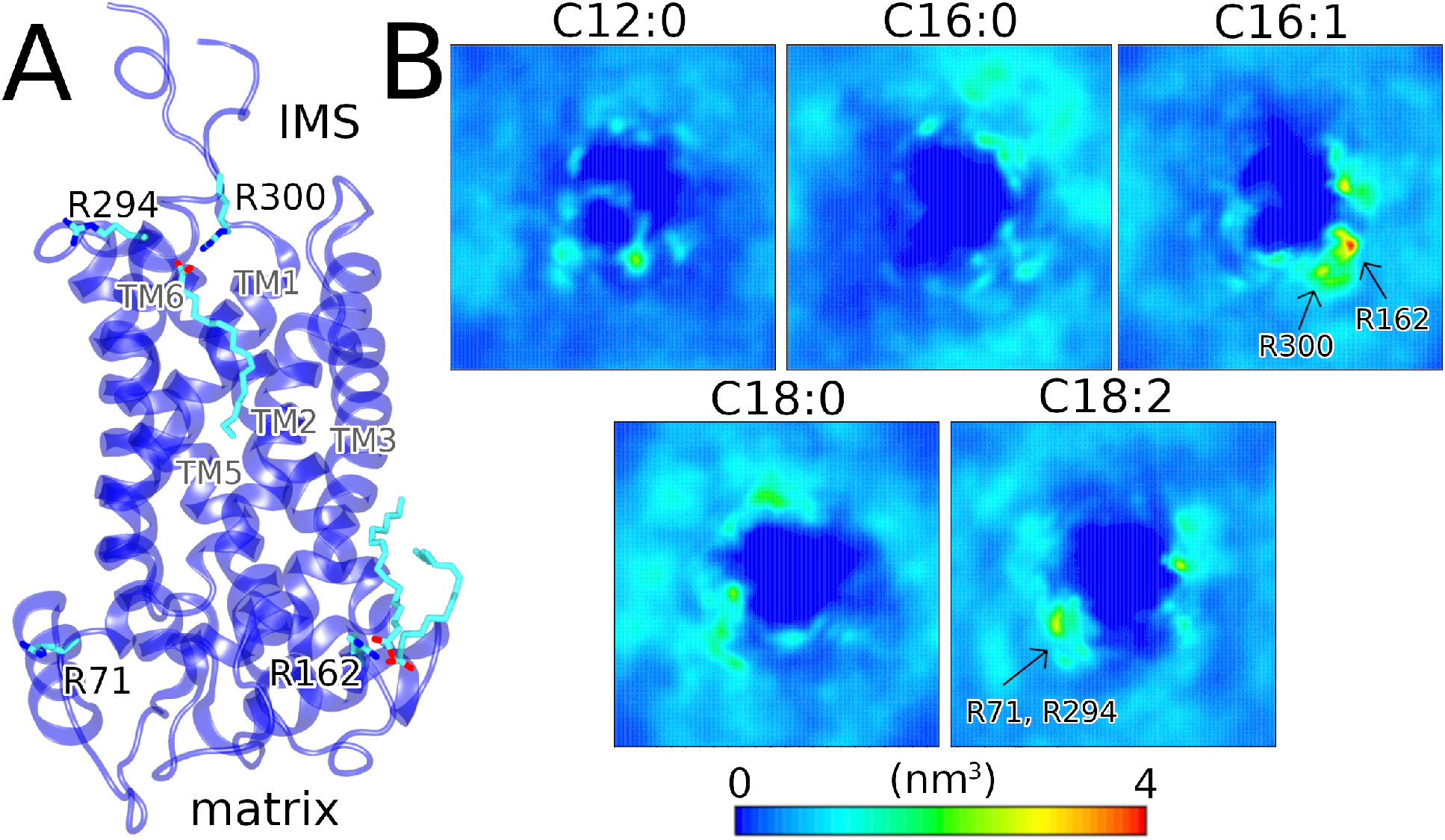
**A**: UCP1 (blue cartoon) with Arginine residues (R71, R162, R294, and R300) that interact with anionic C16:1 fatty acids. Three anionic C16:1 fatty acids are shown binding to UCP1. **B**: 2-dimensional number density of anionic fatty acids in the membrane plane with C12:0, C16:0, C16:1, C18:0, or C18:2 acids. Arrows indicate the amino acid residue hotspots with which the fatty acids interact. Calculations are averaged over all 10 replicas of each system.

Protonatable fatty acids in the translocation pathway is a collective prerequisite for H^+^ leak through UCP1 and AAC. However, the binding site of anionic fatty acid appears different. Whereas fatty acids act as transport substrates in UCP1, fatty acids act as co-factors in AAC.^48^ For AAC, R79 (R84 in UCP1) was proposed to be a fatty acid binding site from molecular dynamics simulations. ^49^ In a more detailed investigation with longer timescale simulations, K22, R79, and R279 (D28, R84, and R277 in UCP1), which otherwise bind the nucleotide, were found to bind the carboxylate group of fatty acids, ^50^ while the fatty acid tails occupied an aperture between TM5 and TM6. An additional fatty acid was found entirely docked into the nucleotide binding site.

The 2-dimensional density maps of anionic fatty acids (Fig. 7) reveal smeared out distributions inside UCP1 in the case of lauric (C12:0) and palmitoleic (C16:1) acid. The central densities are due to fatty acids in the bilayer leaflet facing the IMS. Lauric acid (C12:0) enters UCP1 from the gap between TM5 and TM6 in one replica, and between TM1 and TM2 in another replica. Palmitoleic acid (C16:1) also enters through the gap between TM1 and TM2. The data shows that it is possible for fatty acids to enter the central cavity of UCP1 from the IMM, as recently shown in simulations of AAC. ^50^ Our simulations, together with existing data^32,33,46,50^ indicate that fatty acids enter the protein cavity from either TM1 or TM6 for both UCP1 and AAC. We postulate that fatty acid entry from these sites is likely to be relevant for the H^+^ transport mechanism in UCP1. Although R71, R162, R294 and R300 are common interaction sites for multiple fatty acid types (Table 2 and Fig. 7), none of them interact with all five fatty acid types that we have simulated. The difference in interaction behaviour across multiple replicas can be a result of the fatty acids being initially distributed differently in the IMM, could imply that different fatty acids types interact in a different manner with UCP1, or that the fatty acid binding affinity is too low for specific and tight binding.

In conclusion, we have used MD simulations and free energy calculations to derive a conformational ensemble of UCP1 which validates a breadth of prior experimental biochemical data about lipid binding, the mechanisms of proton transport, the inhibitory impact of ATP binding and the location of the fatty-acid binding sites. We make the following testable predictions about the mechanism of H^+^ transport and ATP-mediated inhibition. First, we propose that the mechanism by which D28 acts as a key acceptor of a proton in a plausible proton transport pathway through UCP1 also involves R277. In the absence of a H^+^, the negative charge on D28 is compensated by R277. Secondly, binding of ATP switches the protonation states of several matrix-facing Lysine amino acids, which can possibly bind fatty acids during the transport cycle. Third, we demonstrate that anionic (but not neutral) fatty acids can approach from the IMM and bind to UCP1 after entering through the TM1/TM2 or TM5/TM6 interface. However, we did not find a specific fatty-acid binding site which was consistently populated across all fatty acid simulations. The absence of a specific fatty acid binding site can indicate that specific binding is not required at all, as long as the fatty acid can bind to UCP1, and contribute a carboxylate group for H^+^ transport. The lack of convergence to a single fatty acid-binding site could also be a limitation of the limited sampling times of the MD simulations. Surprisingly, the presence of a transmembrane potential does not have a significant impact on the ATP-bound conformations.

A primary limitation of the classical MD method employed in this work is the inability to model proton transfer in an efficient and accurate manner. Although recently developed constant pH methods offer an alternative to obtain accurate ionisation states,^51,52^ they are not useful in the UCP1 context, where the protein is exposed to two compartments of different pH. However, the conformational ensemble developed in this work paves the way for other methods such as reactive MD^53^ and QM/MM methods^54^ to investigate proton transport. The models will also enable simulations of the binding of fatty acids to UCP1 with bound nucleotides. The metadynamics simulations do not completely sample the collective variable (RMSD) despite long simulations (microseconds). One way to improve sampling is to experiment with different hill heights. Another approach will be to prevent ATP escape from the binding site in some replicas by enforcing a repulsive wall-like potential. In the future, simulations of the ATP-bound state with fatty acids will be welcome, because these investigations will delineate the impact of ATP on fatty acid binding, and are likely to provide new insights into the mechanism by which ATP inhibits UCP1-mediated H^+^ transport.

## 3 Methods

The initial UCP1 model was created by AlphaFold.^27,28^ Simulations were performed with Gromacs 2020.5 or newer.^55–57^

### Coarse Grained MD Simulations

Three replicas of coarse grained MD simulation of UCP1 in an IMM were prepared using the MARTINI bilayer maker in CHARMM-GUI. ^58^ The membrane composition was constructed to mimic that of the IMM^59,60^ (Table 3). Cardiolipin parameters for MARTINI3^61^ were not available at the time of setting up the simulations. We used MARTINI2 ^62,63^ with an elastic network model to restrain the tertiary structure of the protein. The solvent was polarizable water with a 0.15 M NaCl concentration. The systems were minimized by 10,000 steps of the steepest descent algorithm and equilibrated during five steps in which restraints on the system were gradually lifted. Each production run was simulated for 3 μs with a 15 fs time step at 1 atm using the semi-isotropic Parrinello-Rahman barostat^64^ with a time constant of 12 ps and compressibility 0.0003/bar. The temperature of the system was kept at 310 K by the velocity rescale thermostat with a time constant of 1 ps. van der Waals and electrostatic interactions were cut off at 1.2 nm. The short-range neighbour list was cut off at 1.2 nm and was updated every 20 step.

### All Atom MD Simulations

All all-atom simulations were prepared using the membrane builder function in CHARMM-GUI. ^58^ The membrane composition was the same as for the coarse-grained simulations (Table 3) except in the simulations with fatty acids where also 5% neutral and 5% anionic fatty acids were added to each leaflet. The simulation systems contained ~28,000 water molecules and had a 0.15 M NaCl concentration. The systems were minimized by 5,000 steps of the steepest descent algorithm and equilibrated during five steps in which restraints on the system were gradually lifted. All all-atom production simulations were run for 500 ns with a 2 fs time step at 1 atm using the semi-isotropic Parrinello-Rahman barostat ^64^ with a time constant of 5 ps and compressibility 0.00045/bar. The temperature was 310 K and was kept by the Nosé-Hoover thermostat^65,66^ with a time constant of 1 ps. The Linear Constraint Solver (LINCS) ^67^ algorithm was used to constrain all bonds containing hydrogen. The cut off for short range interactions and the neighbour list was 1.2 nm and particle mesh Ewald^68,69^ was applied for long range electrostatic interactions. In the simulations with fatty acids an electric field corresponding to a transmembrane electric potentials≃160 mV was applied across the membrane pointing from the IMS toward the matrix. Solvent cavities in the UCP1 model and experimental UCP2 structure were determined with the HOLE program. ^26^ MDAnalysis^70^ and PyLipid^71^ were used to analyse ATP binding conformations.

### Metadynamics Simulations

The Multiple Walker Well Tempered Metadynamics part of the simulations^35,36^ were performed with the COLVARS module extension for GROMACS. ^72^ In the metadynamics simulations without a transmembrane electric potential Gaussian hills were deposited every 1000 step (2000 fs), the hill height was 0.01 kJ/mol, the width of the Gaussian hills were 0.026 nm, and the bias temperature was 1500 K. The parameters for the simulations with a transmembrane electric potential were the same as in the simulations without a transmembrane electric potential, except the hill height and bias temperature were increased to 1.0 kJ/mol and 1500 K, respectively, in the hope for faster convergence.

## Supporting information

Supplementary Movie (ATP binding)

## Acknowledgments

LJ and HK are supported by the Lundbeck foundation Ascending Investigator grant number #R344-2020-1023. Simulations were performed on the DeiC National HPC (g.a. DeiC-SDU-N5-000007), and the Novo Nordisk Foundation-funded ROBUST Resource for Biomolecular Simulations #NNF18OC0032608. We acknowledge PRACE for awarding us access to resource Joliot-Curie SKL based in France at GENCI@CEA. We thank Prof. Edmund Kunji and Dr. Hridya VM for useful discussions.

