## Supplementary Information for "The ATP-bound State of the Uncoupling Protein 1 (UCP1) from Molecular Simulations"

### $H^+$ Transport models

Several uncoupling protein 1 (UCP1) proton ($H^+$) transport models have been proposed through the years among which some are illustrated in Fig. S1:

- **Cycling model (Fig. S1A)** fatty acids cycle between the intermembrane space (IMS) and matrix leaflets of the inner mitochondrial membrane (IMM).[1] In one cycle a neutral fatty acid flips from the IMS to the matrix leaflet where it releases its proton to the matrix space. Next, the transport of the then anionic fatty acid back to the IMS leaflet is mediated by UCP1. In the IMS leaflet the fatty acid is again protonated and the cycle repeated.[1–3]

- **Functional competetion model (Fig. S1B)** UCP1 independently transport $H^+$ and the function of fatty acids is to remove the inhibiting purine nucleotide by competitive binding.[2]

- **Shuttling model (Fig. S1C)** a fatty acid binds with its carboxyl group inside the central cavity of UCP1.[4] The carboxyl group shuttles back and forth between the IMS and matrix where it accepts and releases $H^+$, respectively, while the fatty acid chain stays bound.[4]

- **Buffering model (Fig. S1D)** one or two fatty acids bind inside the pore of UCP1 to provide additional carboxyl groups that together with titratable amino acids in UCP1 constitute a complete $H^+$ translocation pathway through UCP1.[2,5]

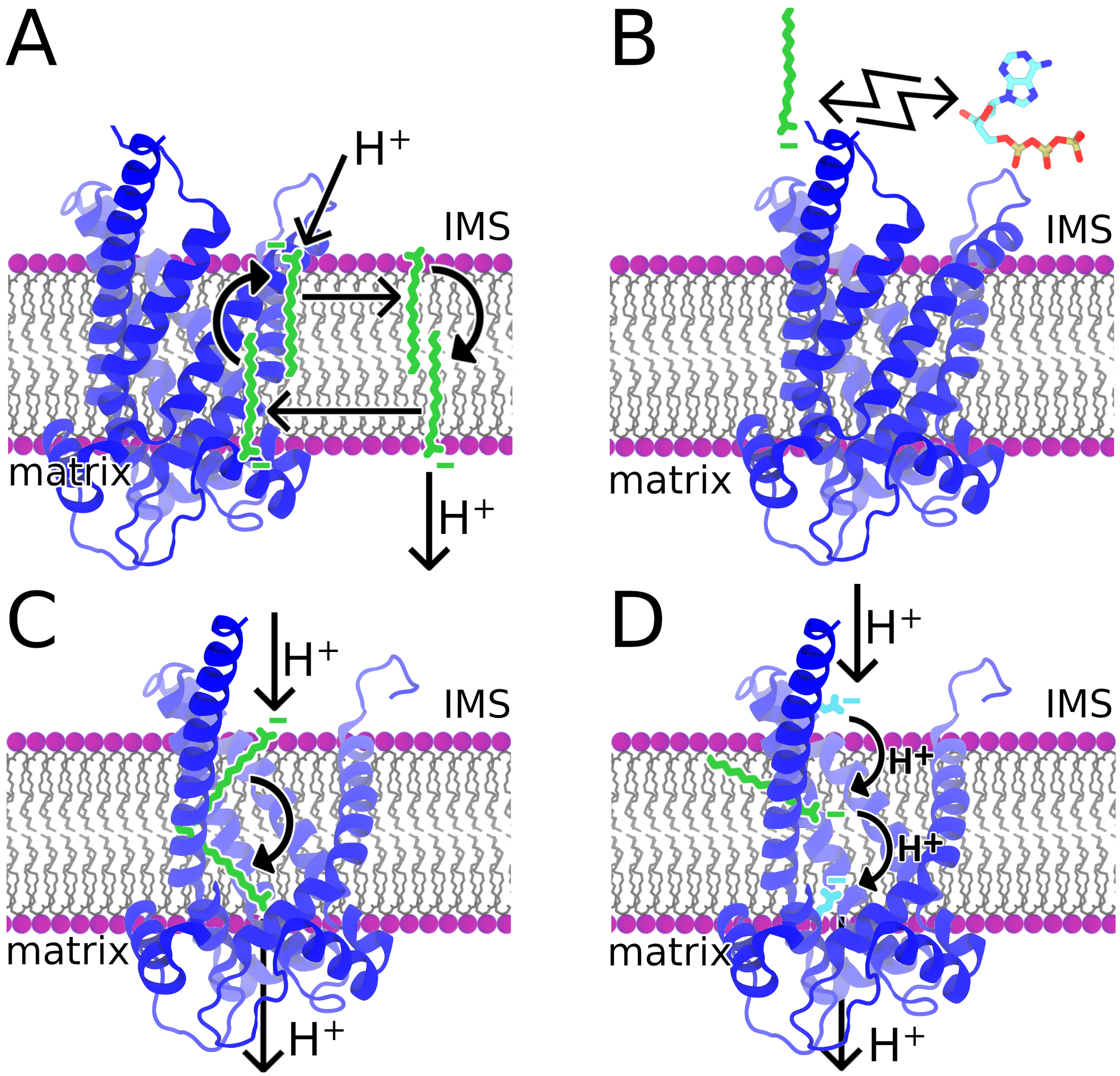


Figure S1: Different $H^+$ transport models. **A**: cycling model, **B**: functional competition model, **C**: shuttling model, and **D**: buffering model.

#### UCP1 Structure

Figure S2 depicts the AlphaFold[6,7] UCP1 model from different perspectives with respect to the IMM. The UCP1 model possesses the three fold symmetry common for proteins in the mitochondrial carrier family.[8] It folds into three repeated domains of ∼100 residues that each contains two transmembrane $\alpha$-helices (H1-H6), which fold into a helix pair, and a matrix facing short $\alpha$-helix (h1-h3) and loop (l1-l3) connecting to the neighbouring repeat.[9]

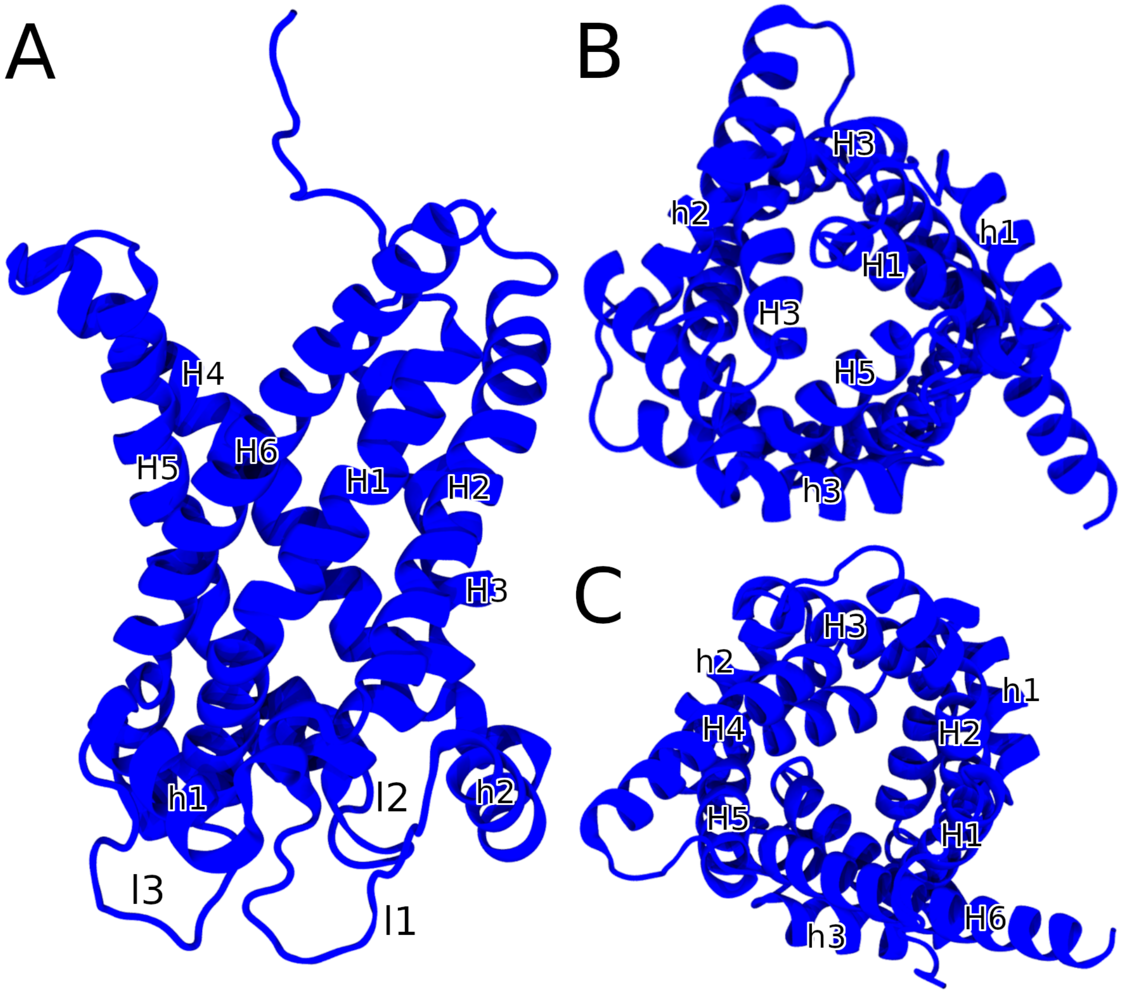


Figure S2: UCP1 AlphaFold model in blue cartoon seen from the IMM (**A**), matrix (**B**), and IMS (**C**). H1-H6 represents the six transmembrane helices, h1-h3 and l1-l3 are the matrix facing $\alpha$-helices and loops that connect the three transmembrane helix pairs, respectively.

### Binding of Cardiolipin to UCP1

Figure S3A shows the membrane densities of cardiolipin phosphate atoms in three coarse grained molecular dynamics simulations of UCP1 in an IMM. In all three replicas the same increased densities can be seen with a three fold symmetry corresponding to cardiolipins at the three predicted binding sites.[10] Cardiolipin spontaneously diffuse to and bind the predicted cardiolipin binding site[10] on UCP1 shown in Fig. S3B. The density of cardiolipin phosphate atoms is higher near residues 55-57 and 268-270 because a cardiolipin was coincidentally placed nearby the site initially.

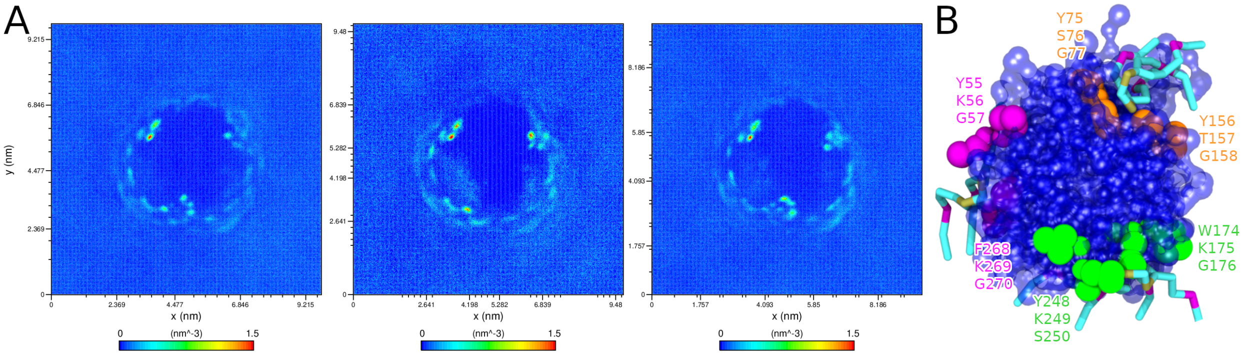


Figure S3: **A**: 2D membrane density maps of cardiolipin phosphate atoms in three 3 µs replicas of coarse grained molecular dynamics simulations of UCP1 in an IMM. The density maps are calculated based on the last 2 µs of the simulations. **B**: Binding of three cardiolipins that spontaneously diffuse to and bind the predicted cardiolipin binding sites on UCP1[10] indicated by coloured spheres.

Figures S4-S5 show the radial distribution functions (RDF) of the phosphate atoms of the three bound cardiolipins with respect to UCP1 in simulations of UCP1 with ATP in the binding site cavity with (E191$^{-}$) and (E191p), respectively. Figure S6 show the membrane density maps of cardiolipin in simulations of UCP1 in an IMM with different fatty acids. In all simulations the three cardiolipins remain bound to UCP1.

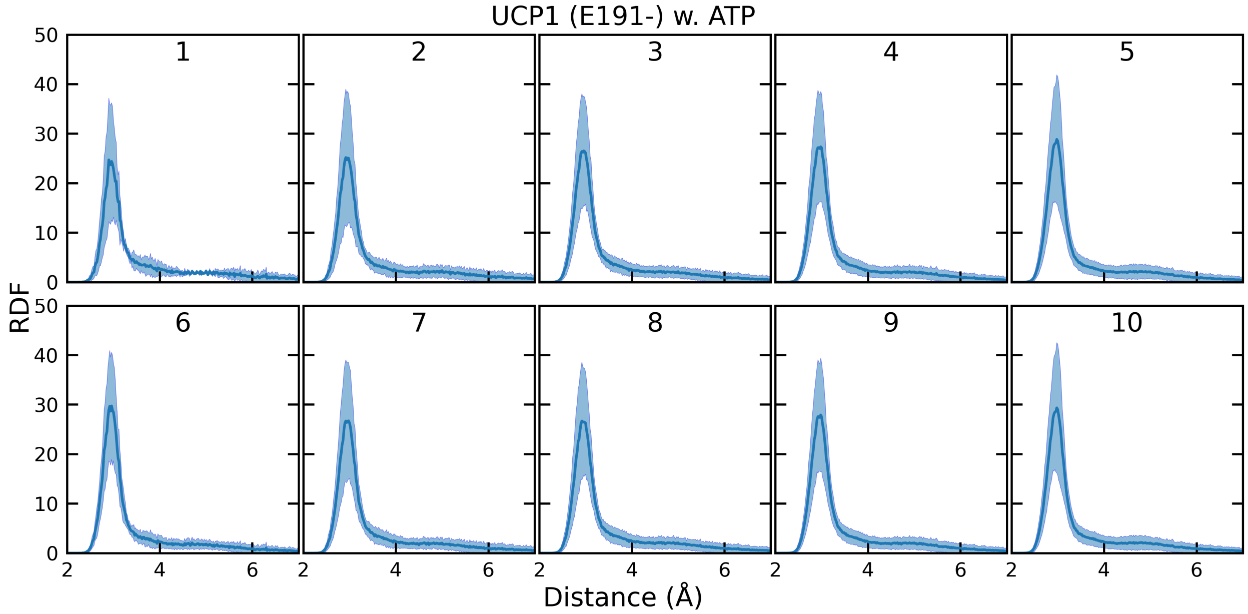


Figure S4: RDF of cardiolipin phosphate atoms with respect to UCP1 in simulations of UCP1 (E191$^{-}$) in an IMM with ATP placed in the mouth of UCP1.

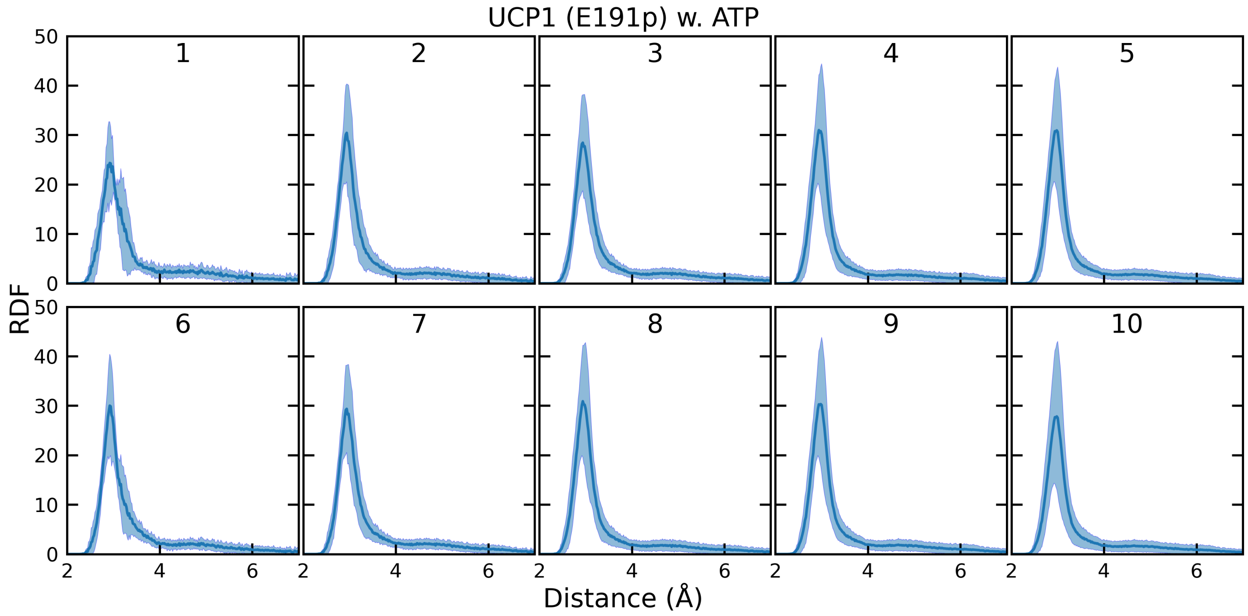


Figure S5: RDF of cardiolipin phosphate atoms with respect to UCP1 in simulations of UCP1 (E191p) in an IMM with ATP placed in the mouth of UCP1.

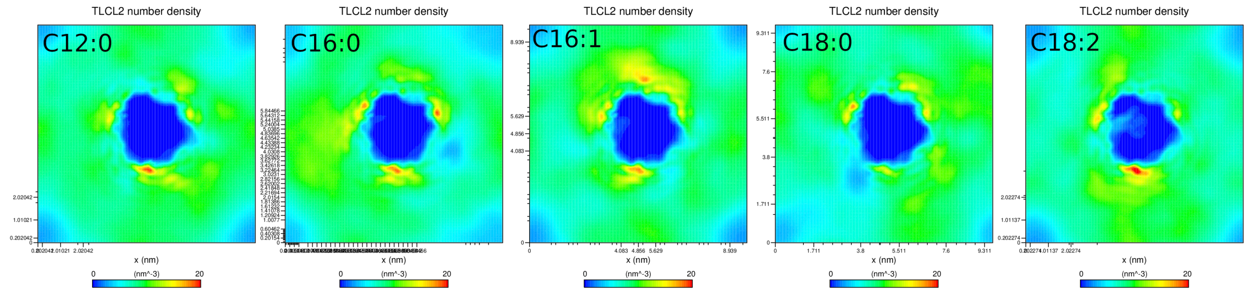


Figure S6: 2D membrane density maps of cardiolipin averaged across all 10 simulation replicas of UCP1 in an IMM with lauric (C12:0), palmitic (C16:0), palmitoleic (C16:1), stearic (C18:0), and linoleic (C18:2) acid.

### RMSD

Figures S7-S12 show the root mean square displacement (RMSD) as a function of simulation time in the all atom simulations.

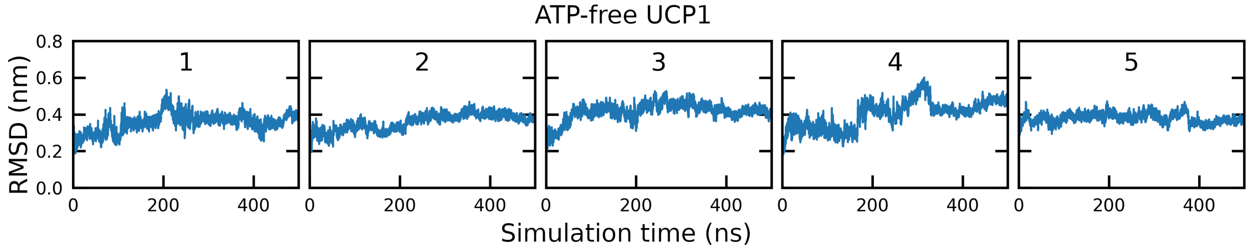


Figure S7: RMSD of the protein backbone in simulations of ATP-free UCP1 ($E191^{-}$) in an IMM.

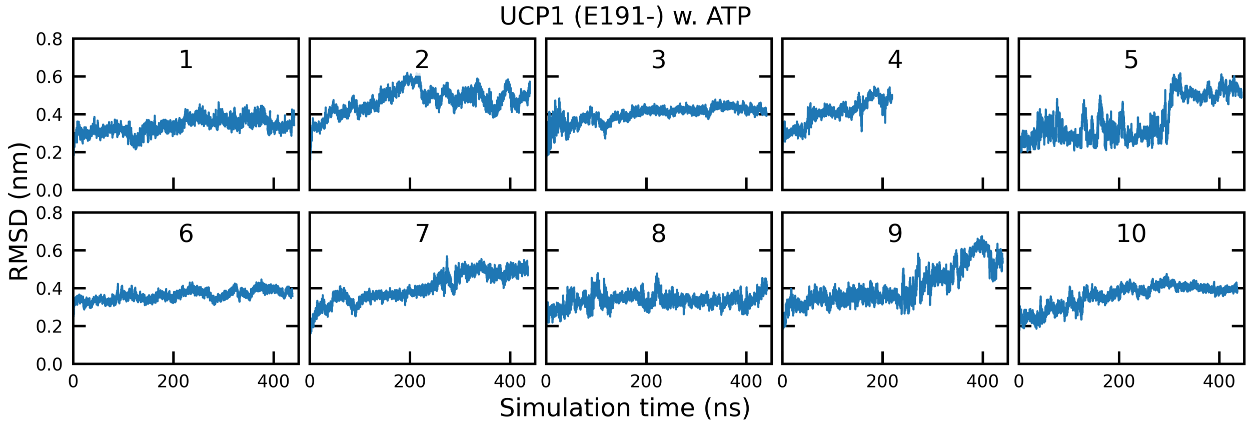


Figure S8: RMSD of the protein backbone in simulations of UCP1 ($E191^{-}$) with ATP in an IMM. Simulation 4 was cut short because ATP drifted away from UCP1.

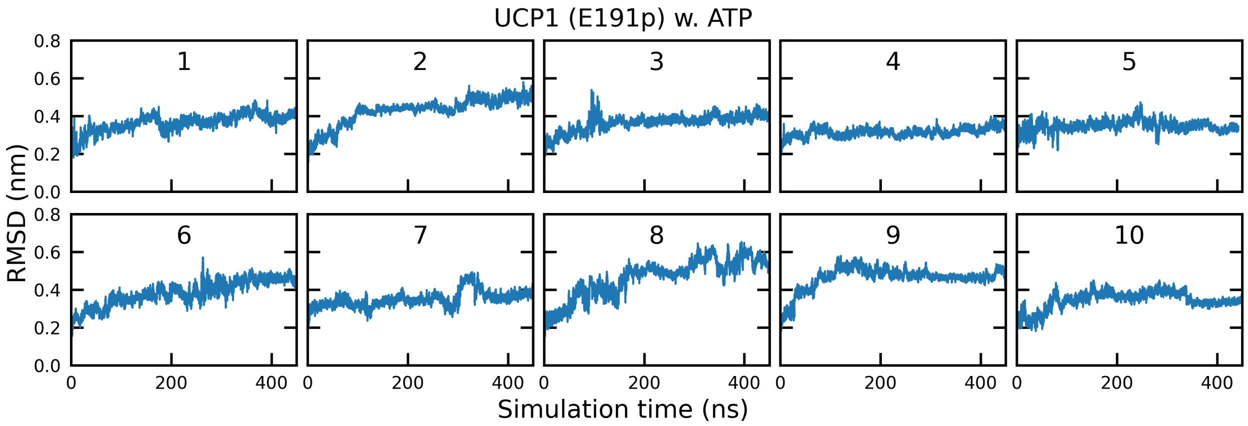


Figure S9: RMSD of the protein backbone in simulations of UCP1 (E191p) with ATP in an IMM.

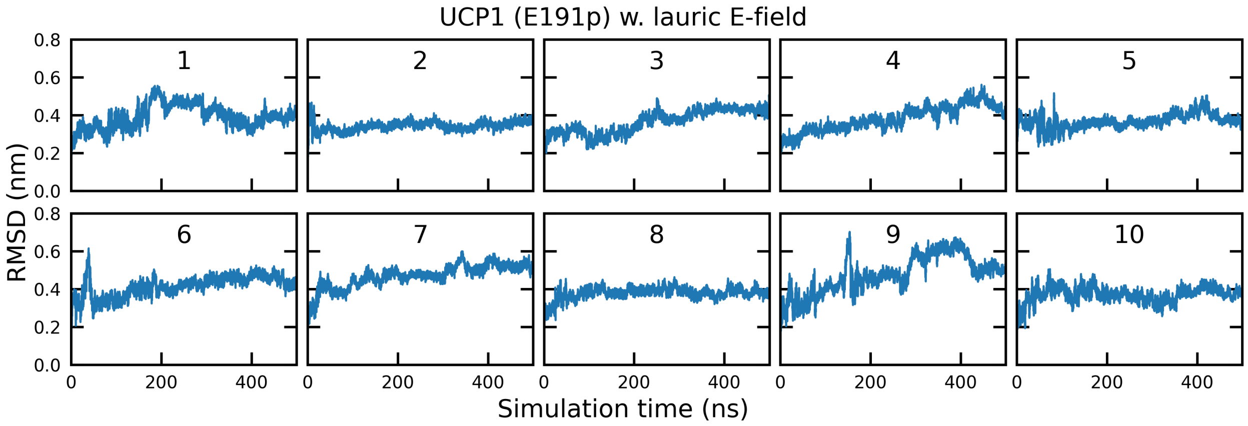


Figure S10: RMSD of the protein backbone in simulations of UCP1 (E191p) in an IMM with lauric acid and an external electric field.

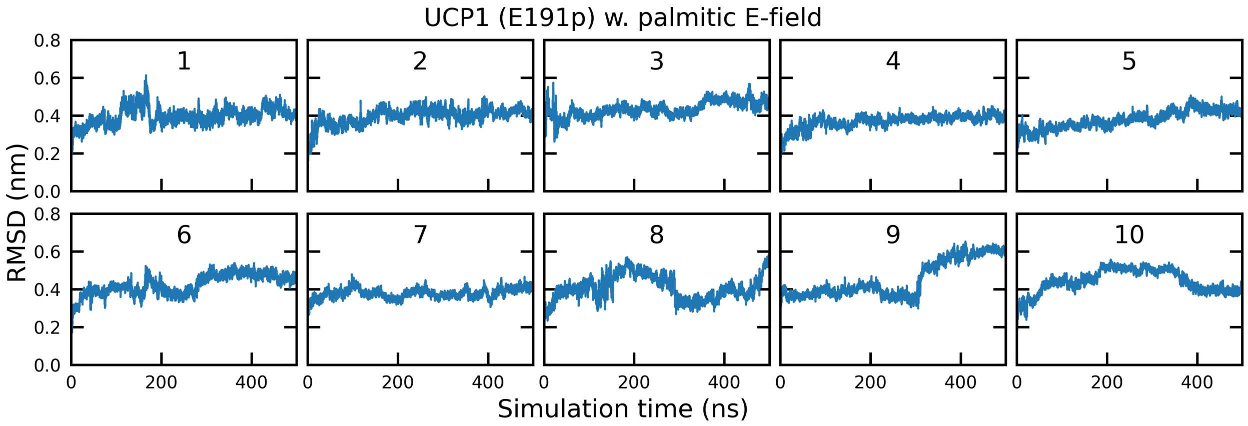


Figure S11: RMSD of the protein backbone in simulations of UCP1 (E191p) in an IMM with palmitic acid and an external electric field.

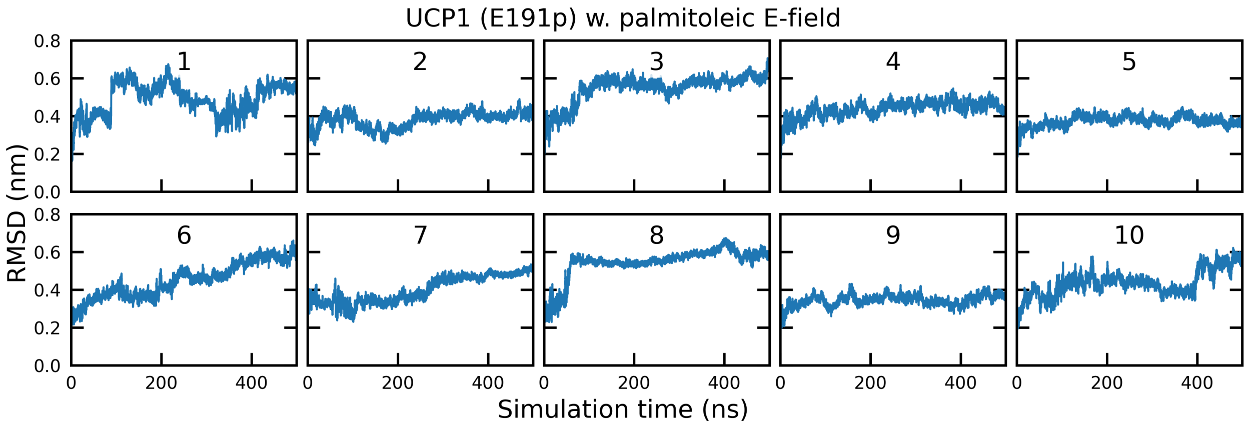


Figure S12: RMSD of the protein backbone in simulations of UCP1 (E191p) in and IMM with palmitoleic acid and an external electric field.

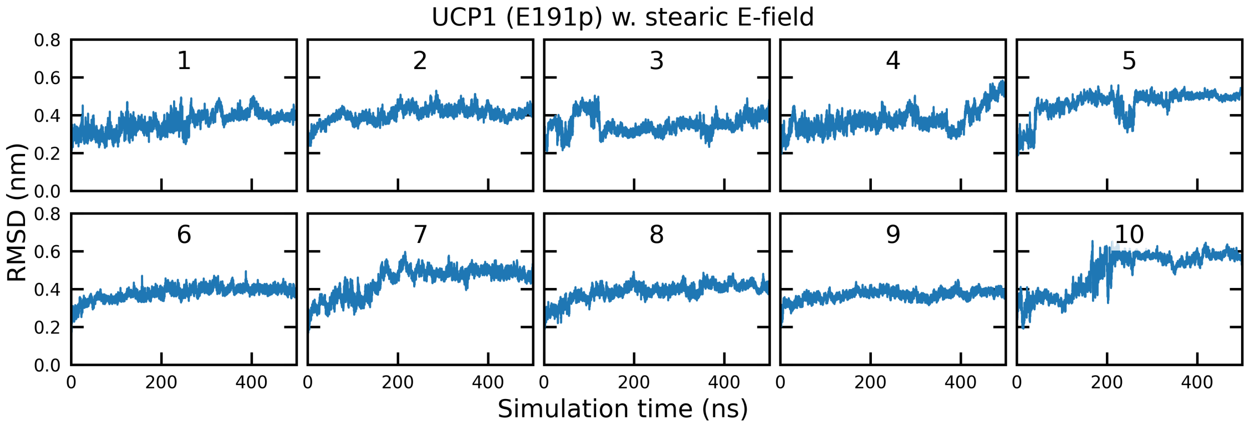


Figure S13: RMSD of the protein backbone in simulations of UCP1 (E191p) in an IMM with stearic acid and an external electric field.

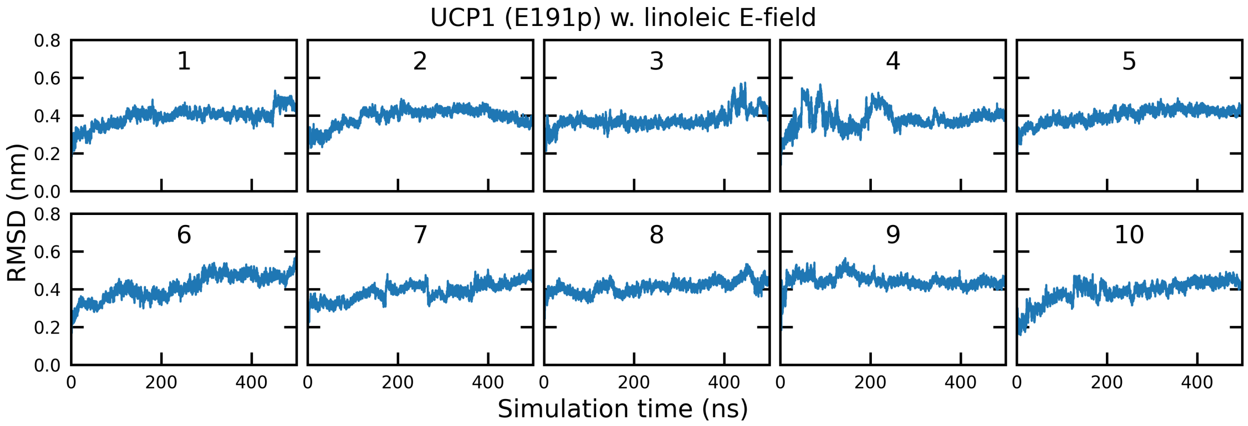


Figure S14: RMSD of the protein backbone in simulations of UCP1 (E191p) in an IMM with linoleic acid and an external electric field.

### Initial and Final ATP Binding Configurations

Figure S15 illustrates the initial (green) and final (red) binding configuration of ATP in simulations with E191$^-$. Five simulations (1-5) were prepared with the nucleobase facing the purine nucleotide binding site and five simulations (6-10) were prepared with the triphosphate group facing the purine nucleotide binding site. In none of the ten simulations ATP entered the binding site (three central orange Arginine residues) mainly because the R92-E191 salt bridge obstructed entrance to the binding site.

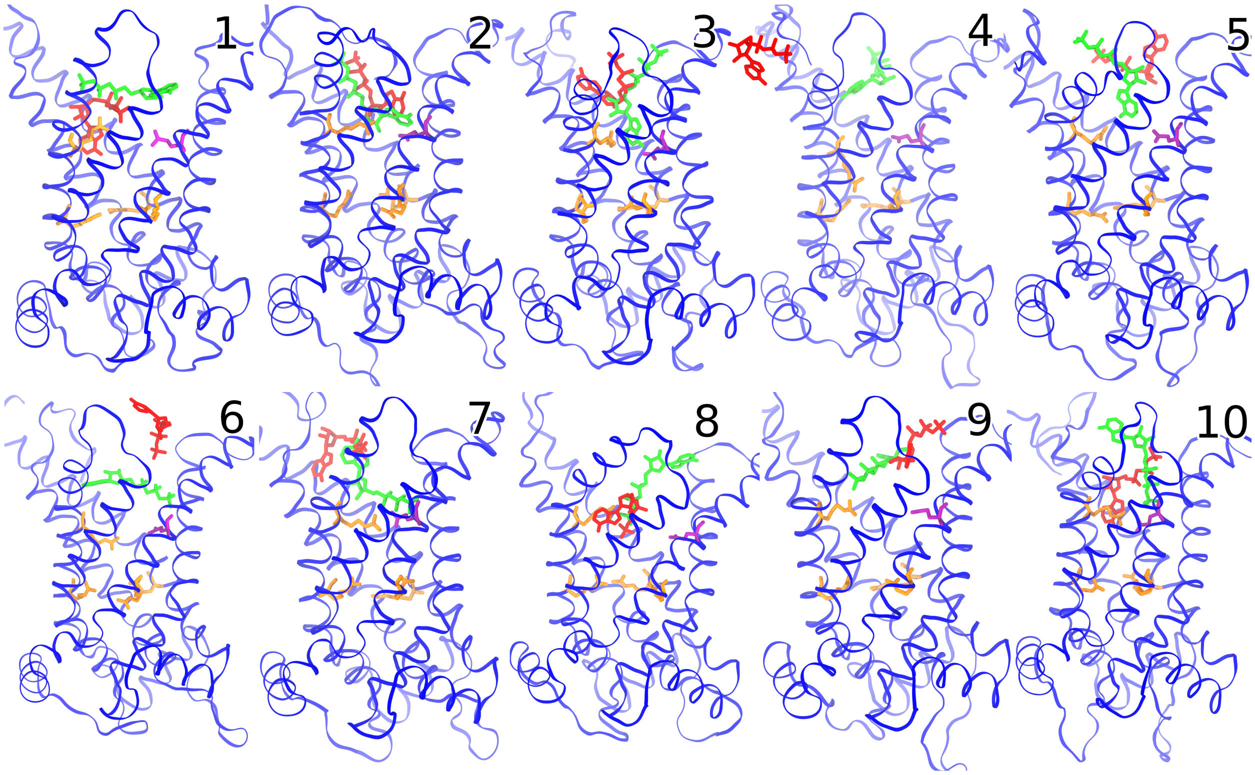


Figure S15: The initial (green sticks) and final (red sticks) binding configuration of ATP in the simulations of UCP1 (E191$^-$) with ATP. The purine nucleotide binding site is marked by the central orange arginine residues. The salt bridge gating the the purine nucleotide binding site is highlighted in purple (E191) and orange (R92).

### R92 Shuttle

R92 binds ATP from the IMS and shuttles it to the binding site where R92 continuous to bind and stabilize the ATP binding configuration. Figure S16 shows the centre of mass distance between R92 and ATP as a function of simulation time.

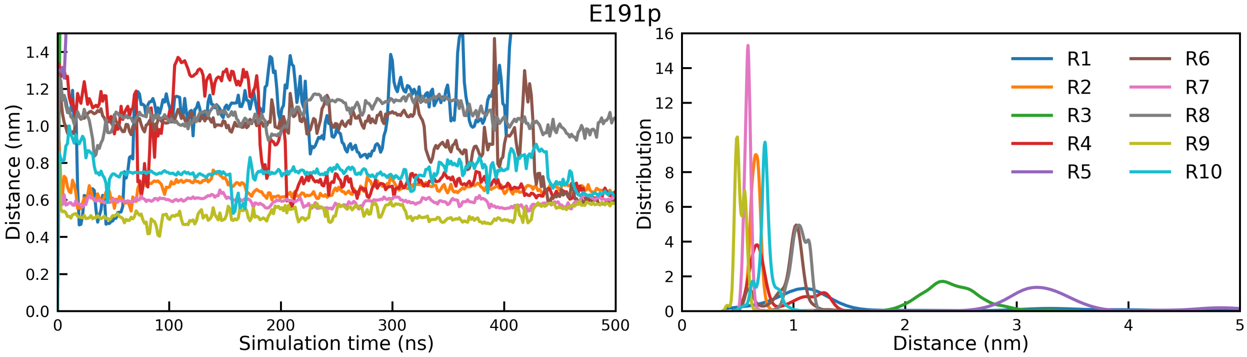


Figure S16: Center of mass distance between R92 and ATP during the ten simulations of UCP1 with ATP as a function of simulation time (left) and the associated distributions (right).

#### R277-D28 Distance

R277 and D28 bind to each other. Figure S17 shows the distance between R277 and D28 as a function of simulations time.

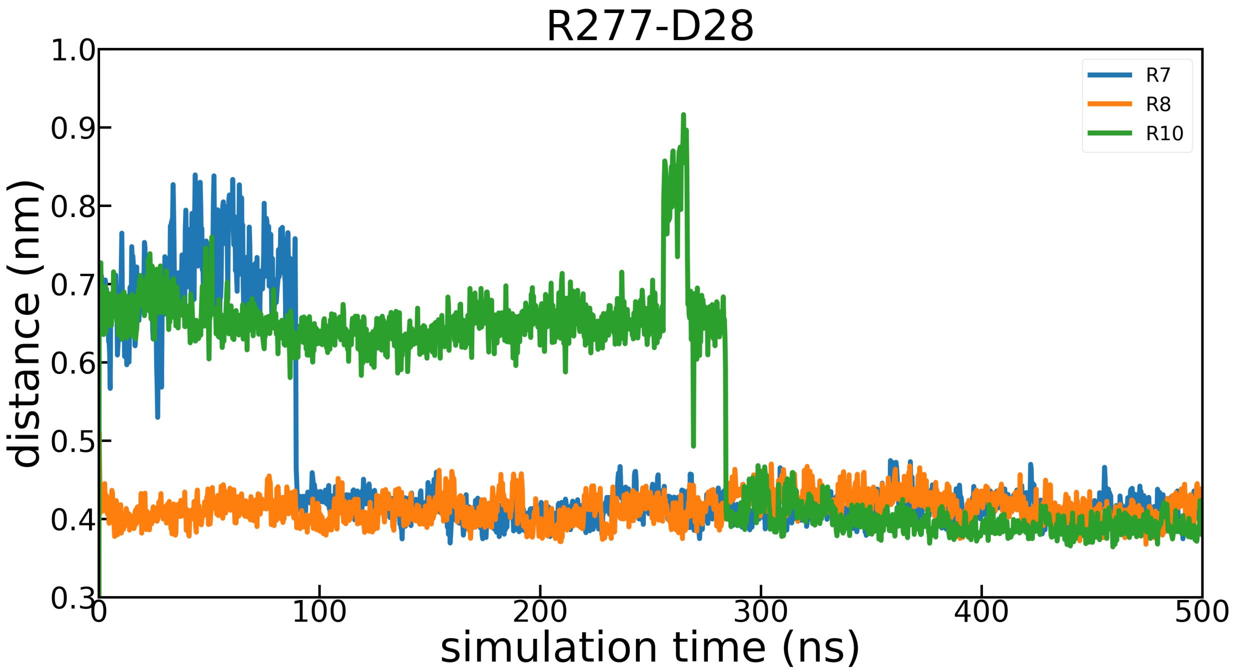


Figure S17: Distance between the R277 guanidium group carbon atom and the D28 carboxyl group carbon atom, during simulation replicas R7, R8, and R10 with E191p.

### pKa Values of Titratable Amino Acids

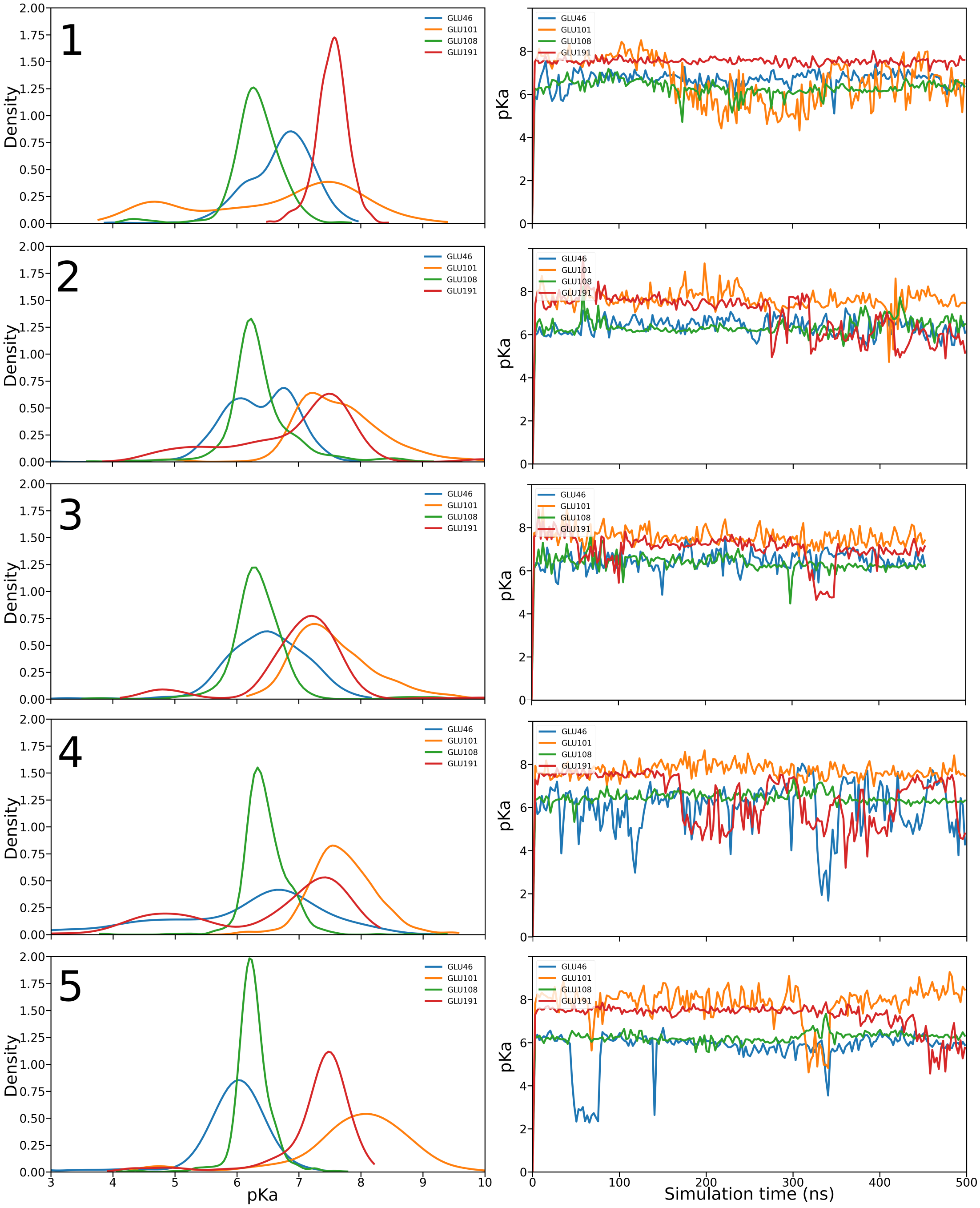


Figure S18: Distribution and time evolution of pKa values of all Glumatic Acid residues which had average pKa values above 5.5 in simulations of ATP-free UCP1 in an IMM.

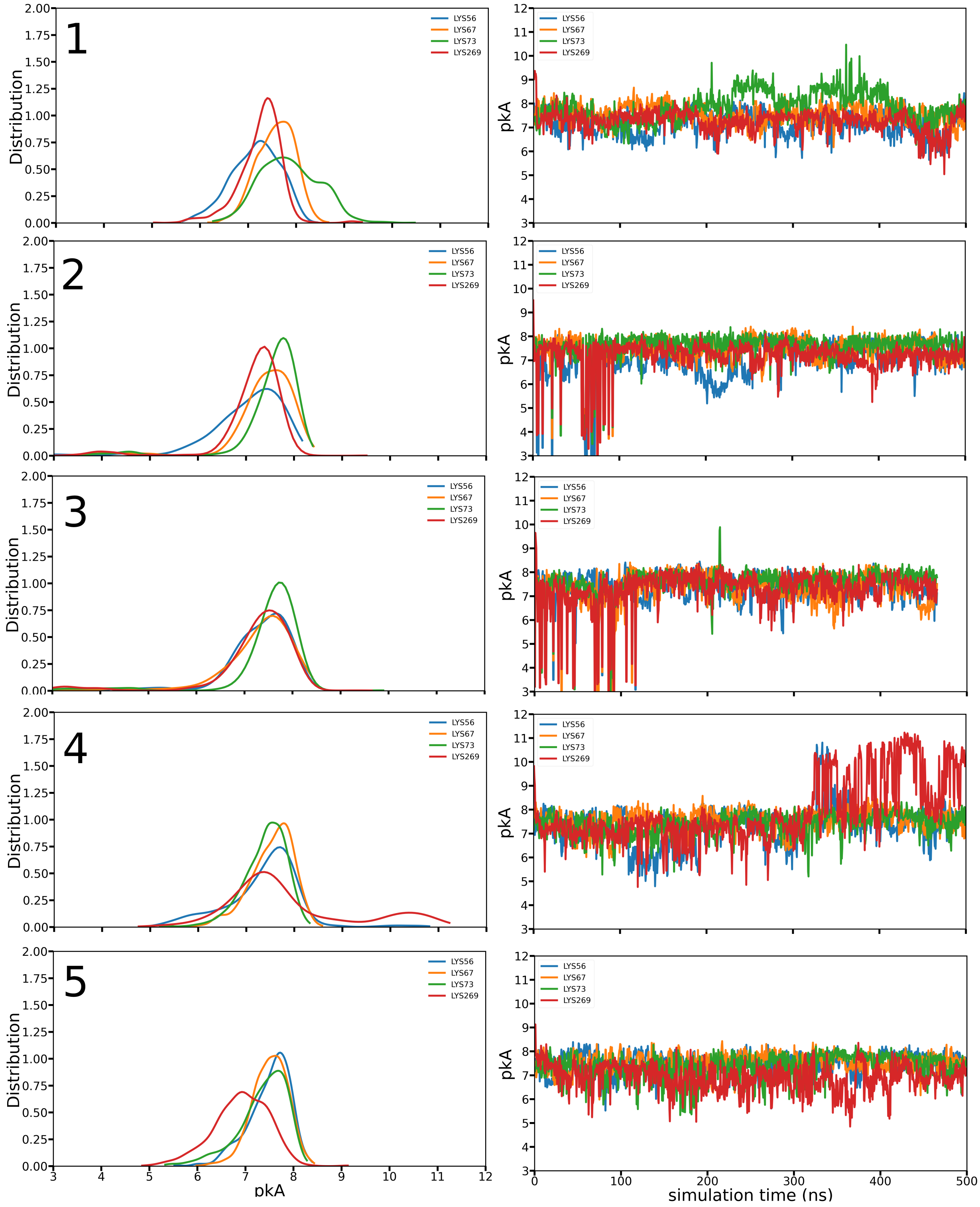


Figure S19: Distribution and time evolution of pKa values of all Lysine residues which had average pKa values below 9 in simulations of unbound UCP1 in an IMM.

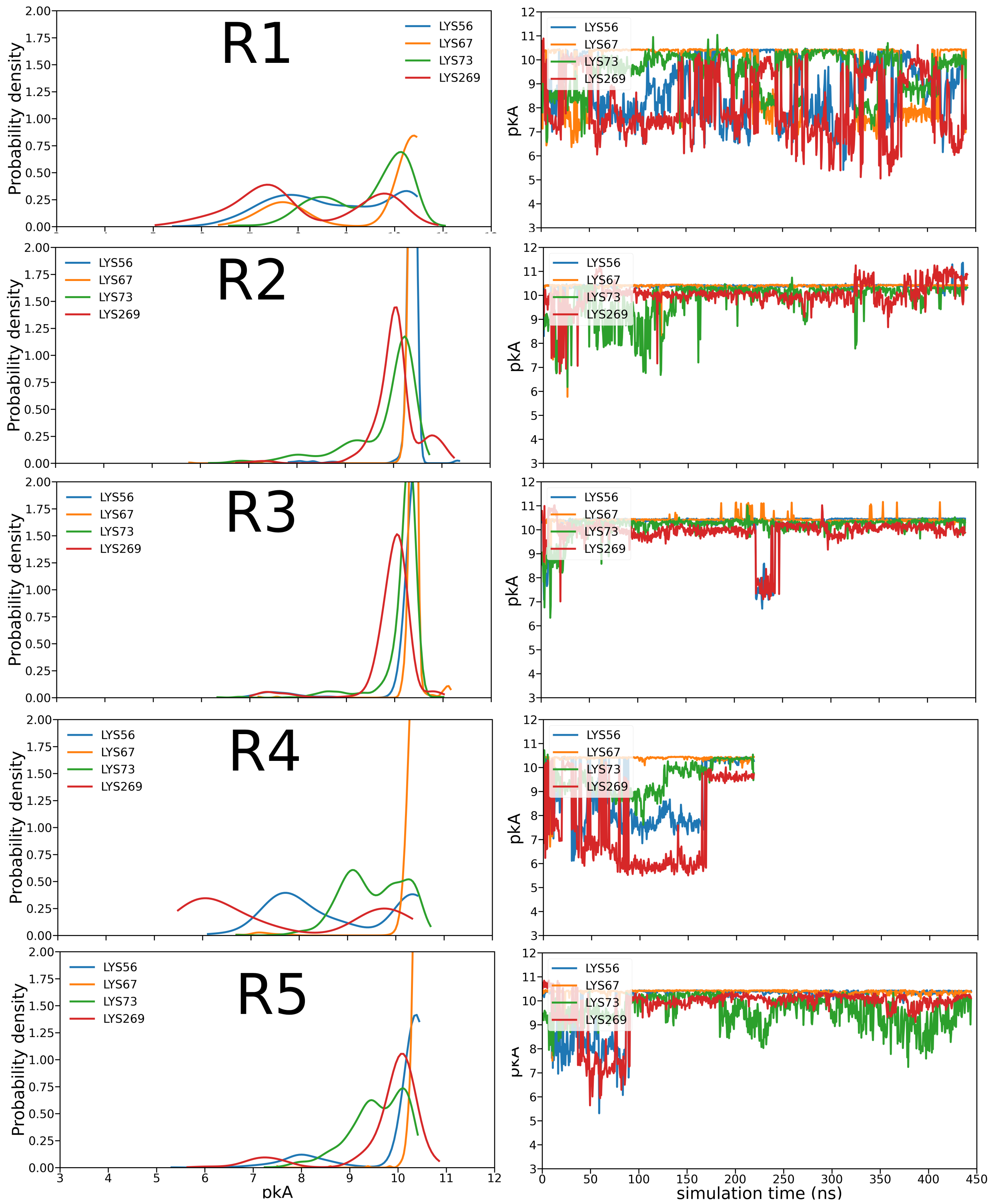


Figure S20: Distribution and time evolution of pKa values of K56, K67, K73, K269 in simulations of UCP1 (E191$^{-}$) in the presence of ATP. Simulation R4 is only ~250 ns because the ATP molecule had drifted away from UCP1 by then. Continued in S21.

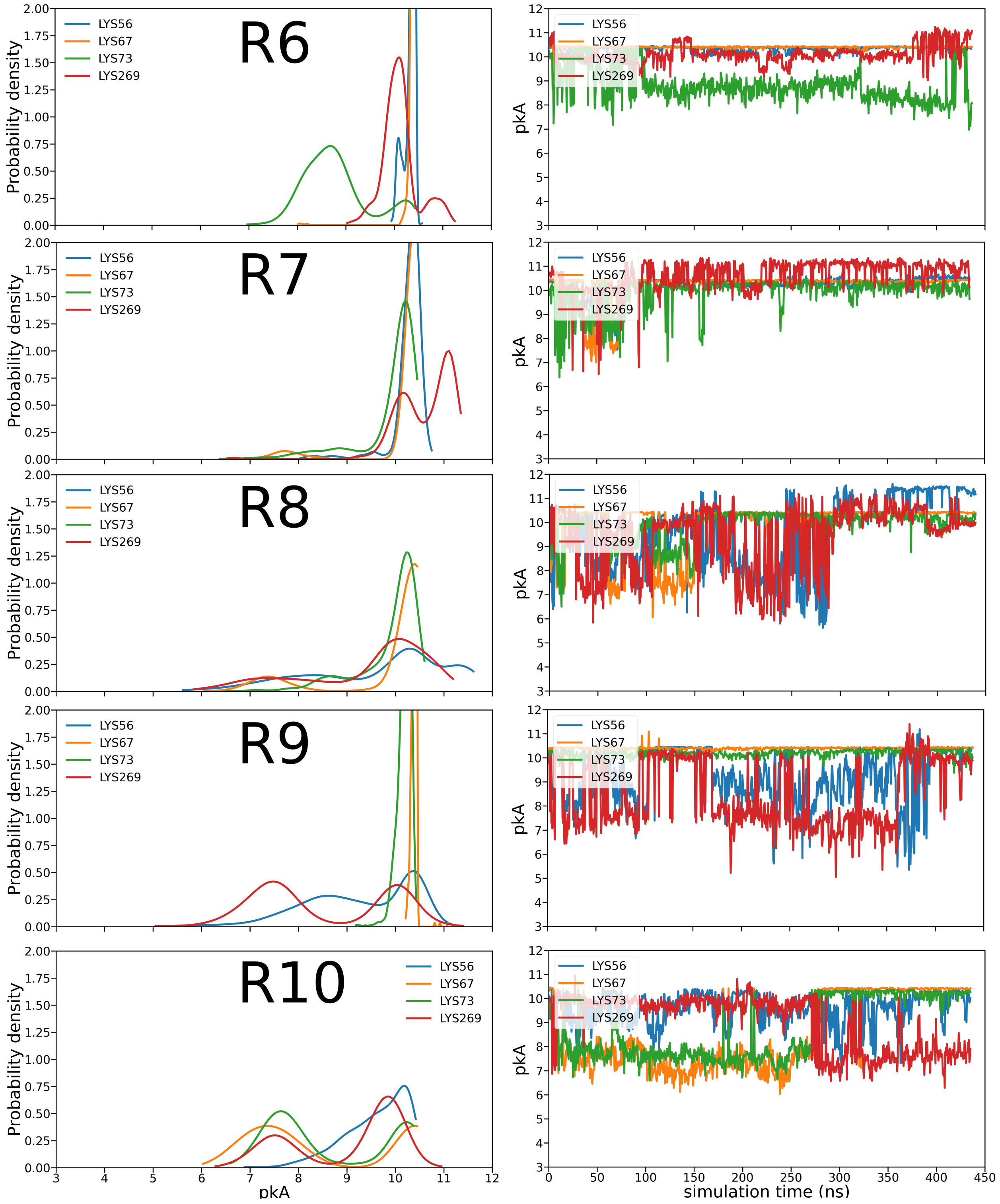


Figure S21: Continuation of Fig. S20. R1 through R10 represent the 10 replicas.

### Neutral Fatty Acids Density Maps

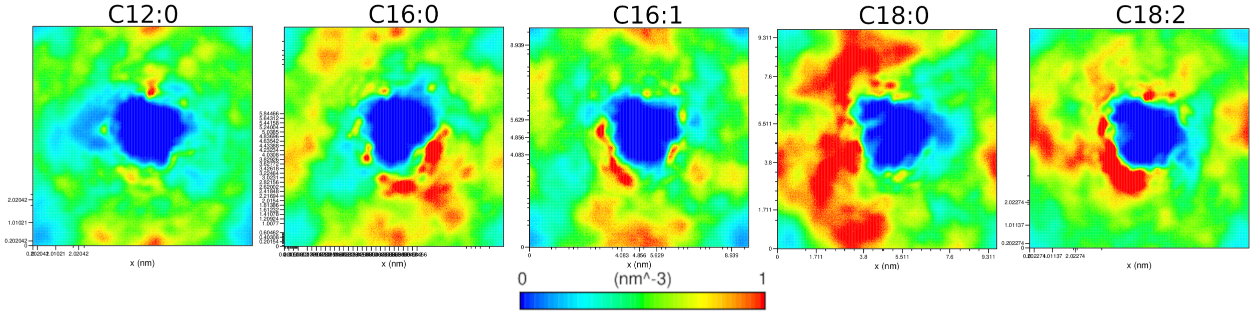


Figure S22: 2D membrane density maps of neutral fatty acids averaged over all 10 simulations of UCP1 in an IMM. Notice that the scale goes from 0-1 $nm^3$ as compare to 0-4 $nm^3$ in the density maps with anionic fatty acids in the main paper.

### Metadynamics Energy Landscapes

Figures S23 and S24 show the energy landscapes generated from multiple walker well tempered metadynamics simulations of ATP bound UCP1 without and with a membrane potential, respectively.

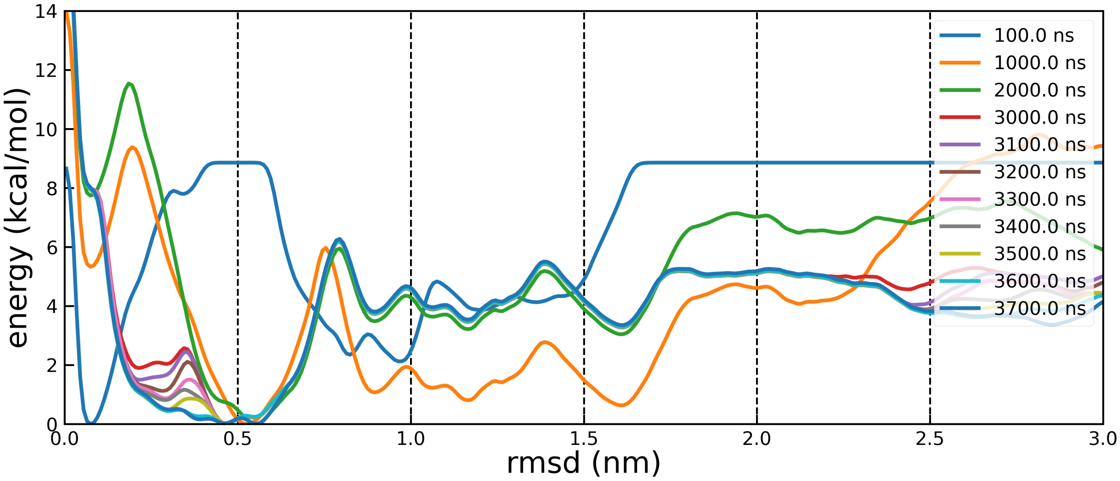


Figure S23: Energy landscapes at different simulation times during metadynamics simulations of UCP1 with bound ATP and no membrane potential.

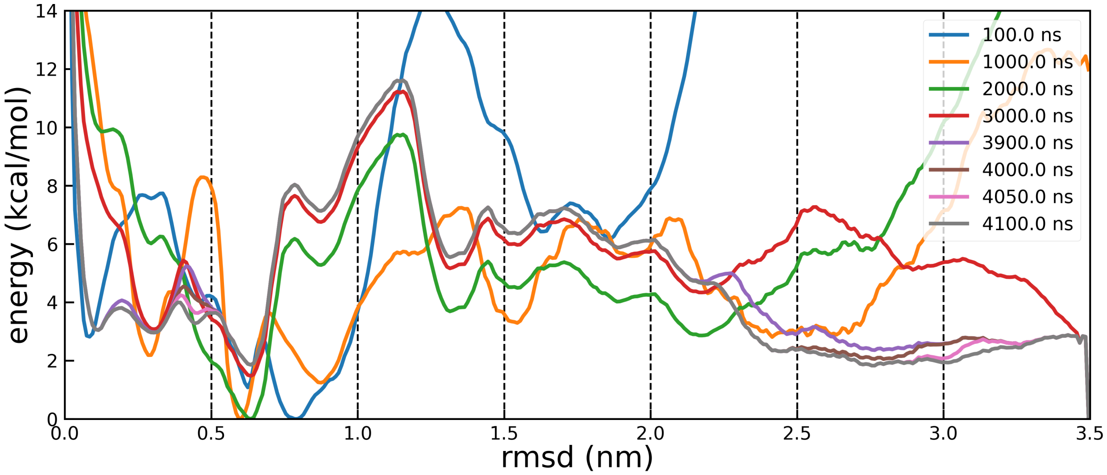


Figure S24: Energy landscapes at different simulation times during metadynamics simulations of UCP1 with bound ATP and a membrane potential. The steep decline at 3.5 nm after 3900 ns is an artefact of the RMSD reaching the predefine boundary of the collective variable and does not represent an energy minimum.

(7) Varadi, M.; Anyango, S.; Deshpande, M.; Nair, S.; Natassia, C.; Yordanova, G.; Yuan, D.; Stroe, O.; Wood, G.; Laydon, A.; Žídek, A.; Green, T.; Tunyasuvunakool, K.; Petersen, S.; Jumper, J.; Clancy, E.; Green, R.; Vora, A.; Lutfi, M.; Figurnov, M.; Cowie, A.; Hobbs, N.; Kohli, P.; Kleywegt, G.; Birney, E.; Hassabis, D.; Velankar, S. AlphaFold Protein Structure Database: massively expanding the structural coverage

of protein-sequence space with high-accuracy models. *Nucleic Acids Res.* **2022**, *50*, D439–D444.

(8) Palmieri, F. The mitochondrial transporter family SLC25: Identification, properties and physiopathology. *Molecular Aspects of Medicine* **2013**, *34*, 465–484.

(9) Crichton, P. G.; Lee, Y.; Kunji, E. R. S. The molecular features of uncoupling protein 1 support a conventional mitochondrial carrier-like mechanism. *Biochimie* **2017**, *134*, 35–50.

(10) Lee, Y.; Willers, C.; Kunji, E. R. S.; Crichton, P. G. Uncoupling protein 1 binds one nucleotide per monomer and is stabilized by tightly bound cardiolipin. *Proc. Natl. Acad. Sci. U. S. A.* **2015**, *112*, 6973–6978.
